# Selenium reduction of ubiquinone via SQOR suppresses ferroptosis

**DOI:** 10.1101/2023.04.13.535674

**Authors:** Namgyu Lee, Sung Jin Park, Mike Lange, Tenzin Tseyang, Mihir B. Doshi, Tae Yong Kim, Yoseb Song, Dong In Kim, Paul L. Greer, James A. Olzmann, Jessica B. Spinelli, Dohoon Kim

## Abstract

The canonical biological function of selenium is in the production of selenocysteine residues of selenoproteins, and this forms the basis for its role as an essential antioxidant and cytoprotective micronutrient. Here, we demonstrate that selenium, via its metabolic intermediate hydrogen selenide, efficiently donates its electrons to ubiquinone to form ubiquinol in the mitochondria through catalysis by sulfide quinone oxidoreductase (SQOR). Hydrogen selenide is superior to hydrogen sulfide as an electron donor owing to its larger valence shell. We show that this mechanism, independently of selenoprotein production, protects against ferroptosis via ubiquinol production in a manner that depends on xCT mediated selenide formation and SQOR activity. Our findings identify a regulatory mechanism against ferroptosis that implicates SQOR and expands our understanding of selenium in biology.

## Main

Selenium is required in biology to produce the amino acid selenocysteine which in turn is required to produce 25 selenoproteins encoded in the human genome ^1, 2^. As selenoproteins include key antioxidant systems such as glutathione peroxidases and thioredoxin reductases, selenium has been clinically explored in various health contexts as an antioxidant. One such selenoprotein, GPX4, is a key regulator of an iron-dependent form of cell death called ferroptosis, as it reduces lipid peroxidation which is the initiating event ^3, 4^. Based on its antioxidant and antiferroptotic roles, selenium has been explored as a preventative or treatment agent in various disease contexts including stroke and cardiomyopathy^5–7^. Recently, it is emerging that selenium may have biological effects outside of this canonical role ^5, 8, 9^. For example, selenium can induce a transcriptional program to protect against ER-stress induced cell death, neurotoxicity and ferroptosis in cerebral ischemia ^5^, and selenium can result in the selenation of cysteine residues of proteins with functional implications in brown adipose tissue^8^.

### A selenoprotein independent function of selenium

The model for the antiferroptotic activity of selenium, in which it is sequentially metabolized then translationally incorporated to selenoproteins, implies that sufficient time to induce GPX4 protein expression is required before protective effects can be seen (Fig 1a). It was recently established that selenium can induce not only the translation of GPX4, but also induce its transcription ^5^. In agreement, we also observe induction of GPX4 transcripts (Sup Fig 1a), and protein level induction of GPX4 (Sup Fig 1c) at 24 hours of treatment with selenite, a common inorganic form of selenium. Surprisingly, while 2 hr of selenite treatment is insufficient to induce either GPX4 transcripts (Sup Fig 1b) or protein levels (Sup Fig 1d-1g), it was sufficient for selenite to exert a robust protective effect against lipid peroxidation induced by GPX4 inhibition (RSL3) (Fig 1b,1c, Sup Fig 2a,2b). Rescue effects against lipid peroxidation at 2 hr (Sup Fig 2a-2c) and later rescue against ferroptotic death (at 24 hr; Sup Fig 2d-2h), were seen in multiple lines against not only RSL3 but also the similar covalent GPX4 inhibitors ML210 and JKE1674, as well as the structurally/mechanistically distinct inhibitor FIN56.

**Figure 1.**
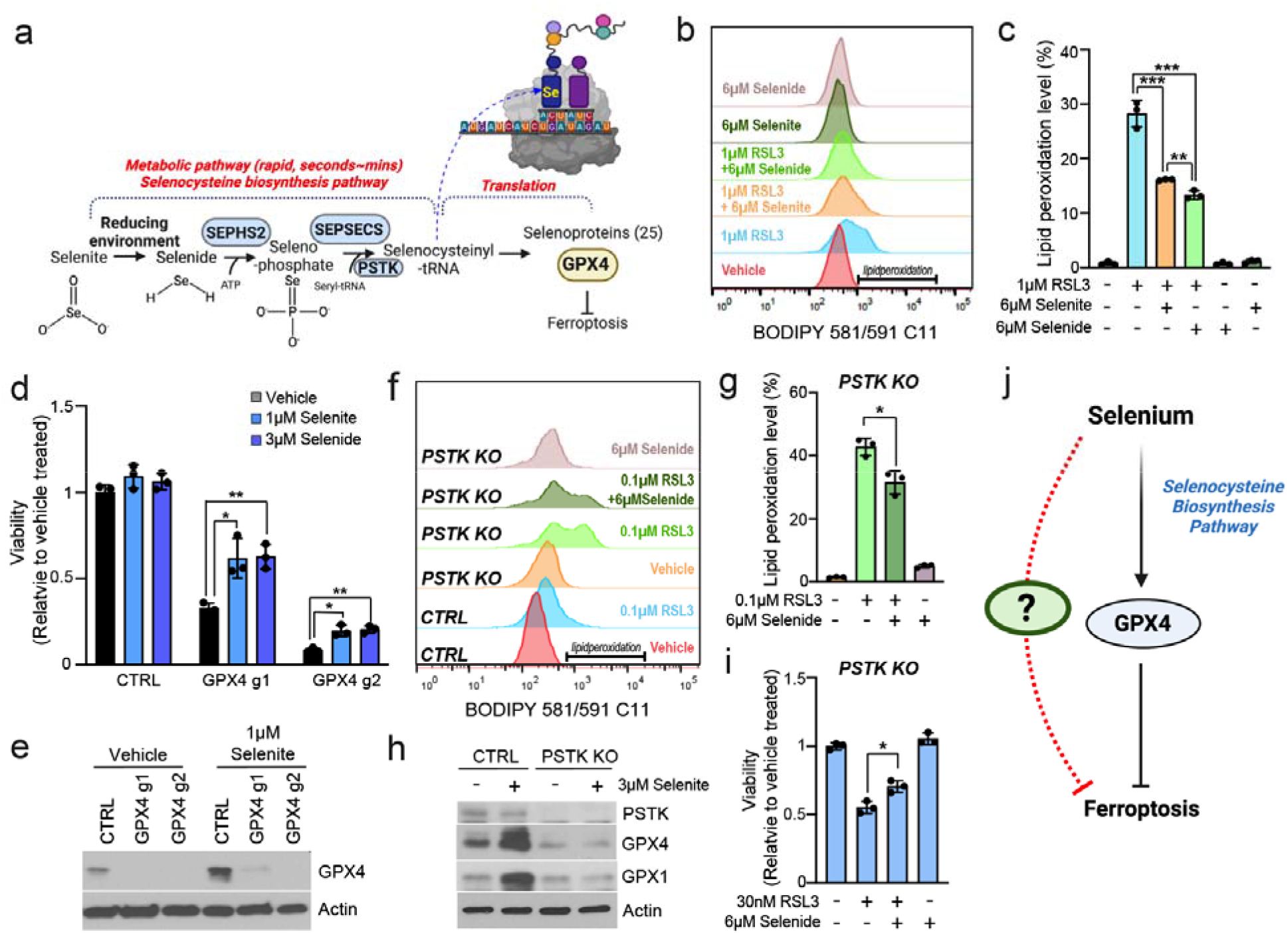
Selenium has an antiferroptotic effect that is independent of selenoprotein production. (a) Schematic diagram of canonical selenocysteine synthesis metabolic pathway and selenoprotein production. (b) BODIPY dye measurement of lipid peroxidation in SK-Hep1 cells treated with 1 µM RSL3 and/or 6 µM selenite or selenide for 1 hr. Bracketed bar indicates the gating for lipid peroxidation. (c) Quantification of lipid peroxidation from in panel b. (d) The viabilities of SK-Hep1 cell line following KO, with gRNAs against GPX4 and vehicle, 1 µM selenite, or 3 µM selenide treatment (blue bars), for 5 d. Values are relative to expression of a non-targeting gRNA in cells vehicle-treated (=1.0). (e) Immunoblots of GPX4 in SK-Hep1 cells treated with 1 µM selenite for 5 d. (f) BODIPY dye measuremen t of lipid peroxidation in CTRL and PSTK KO SK-Hep1 cells treated with 0.1 µM RSL3 and/or 6 µM selenide for 2 hr. Bracketed bar indicates the gating for peroxidized lipid. (g) Quantification of lipid peroxidation in PSTK KO cells from in panel f. (h) Immunoblots of GPX1/4 in CTRL and PSTK KO SK-Hep1 cells treated with vehicle or 3 µM selenite for 24 hr. (i) Viability of SK-Hep1 cells following KO with gRNAs agains t PSTK after treating with vehicle, 30 nM RSL3, with/without 6 µM selenide for 16 hr. (j) Conceptual diagram of our findings suggesting an alternative, rapid mechanism by which selenium can suppres s ferroptosis. Data are mean ± S.D. from biological replicates (*n* = 3) and were analyzed by two-tailed Student’s *t*-test. (c,d,g,i; \**P* < 0.05, \*\**P* < 0.01, \*\*\**P* < 0.001).

In its canonical metabolism to form selenocysteinyl-tRNA, selenite must be converted to selenide, a step in which the xCT transporter has been implicated ^9, 10^. Indeed, we found that directly treating selenide was able to exert the same protective effects (Fig 1b,1c). The protective effect of selenium was even seen under GPX4 KO states (Fig 1d,1e), ruling out induction of GPX4 as the underlying protective mechanism. This observation is in line with previous observations that selenium protection against cell death induced by some insults such as ER stress involve other selenoproteins aside from GPX4 ^5^. To rule out the role of selenium induction of other selenoproteins that could play protective roles, we examined cells devoid of PSTK, a key machinery required in selenocysteinyl-tRNA synthesis (Fig 1a). PSTK KO cells cannot induce selenoproteins in response to selenium (Fig 1h). While these cells were highly sensitive to RSL3 (likely due to limited expression of GPX4), even in this state selenide had a significant protective effect (Fig 1f-1i), demonstrating selenoprotein independent anti­ferroptosis mechanism of selenium.

Collectively, these experiments suggested that selenite has an immediate lipid antioxidant and anti-ferroptotic effect that requires its conversion to selenide but is independent of its canonical function in allowing selenoprotein (GPX4) production (Fig 1j).

### xCT mediates the reduction of selenite to selenide

xCT has been functionally implicated as mediating the step between selenite and selenide in the selenocysteine biosynthesis pathway. The proposed model is that xCT import of cystine ultimately results in its reduction to cysteine and export, providing thiol groups to reduce selenite to selenide ^9, 10^ (Fig 2a). To directly examine this hypothesized model, we utilized our previous lead acetate reactivity method based on sulfur detection chemistry ^9^ (Fig 2b). We further tested selenide detection using a P3 fluorescent probe previously established to detect hydrogen sulfide ^11^, finding that it can also detect selenide (Sup Fig 3a,3b). We observed that various thiol-group-based reducers, including L-cysteine, reduced L-glutathione, β-mercaptoethanol, or N-acetylcysteine could result in selenite to selenide conversion using both the lead acetate colorimetric method and the P3 probe (Fig 2c, Sup Fig 3c,3d). Moving to cells, we observed that xCT activity resulted in thiol accumulation in conditioned media and the thiol levels was correlated with expression of xCT (Fig 2d, Sup Fig 3e-3i,3k), consistent with previous reports^9, 10^. Importantly, xCT inhibitor erastin treatment decreased selenite to selenide conversion in cells as detected by the P3 fluorescence probe (Fig 2e). Erastin also prevented selenium import into cells (Fig 2f), consistent with the notion that selenite to selenide conversion renders it volatile and thus facilitates its import ^9^. This was not a secondary effect from the known role of xCT in glutathione production^12, 13^, as ablating glutathione production via GSS or GCLC did not impact thiol production (Fig 2g, Sup Fig 3i,3j), selenide production (Fig 2h) or selenium uptake levels (Fig 2i). Collectively, these results confirm that xCT activity leads to accumulation of extracellular thiols ^14, 15^, which we demonstrate results in the reduction of selenite to selenide. Thus, while xCT function in glutathione biosynthesis is a key component of cell redox ^12, 13^, our results suggest that xCT’s function in selenide formation also contributes to its well recognized role as a ferroptotic regulator.

**Figure 2.**
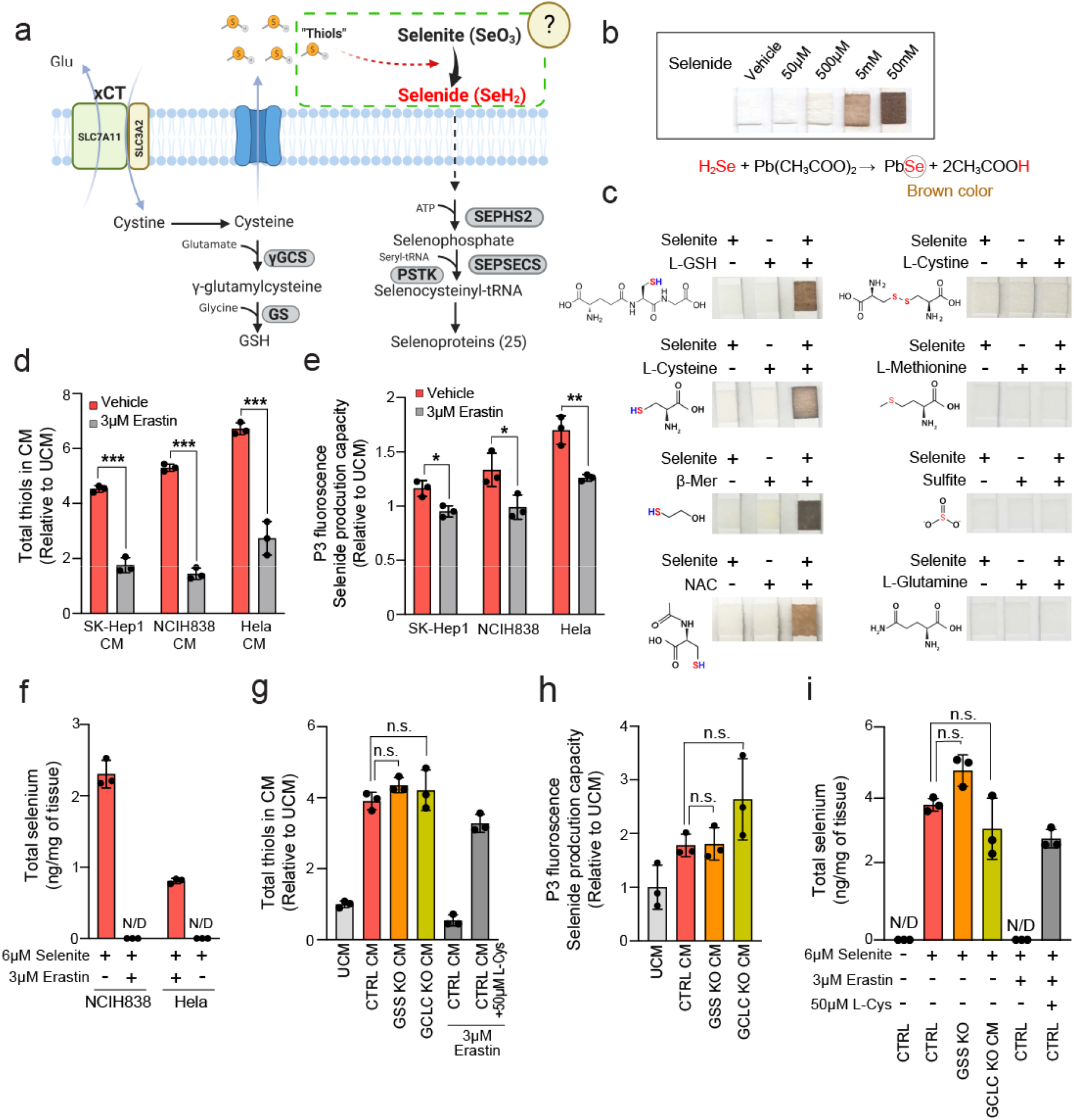
xCT promotes selenide formation independent of GSH production capacity. (a) Schematic diagram of the role of xCT in GSH synthesis and selenoprotein production. The dot square box indicate s the model which has not been experimentally confirmed. (b) Lead acetate embedded paper-based colorimetric selenide detection of different doses of selenide. Selenide results in brown coloration due to reaction with lead acetate. (c) Lead acetate paper after dipping into solutions of selenite and/or different metabolites as indicated. The detection probe was exposed immediately after mixing selenite with thiol-containing metabolites (L-glutathione (L-GSH), L-Cysteine, β-mercaptoethanol (β-Mer), N-acetylcysteine), with negative controls of sulfur non-thiol metabolites (L-Cystine, L-Methionine, Sulfite), or non-sulfur, non-thiol metabolite L-Glutamine. (d) Total thiol measurement of conditioned media from vehicle or 3 µM erastin treated SK-Hep1, NCIH838, and Hela cells, conditioned for 24 hr. Each value i s relative to that of the unconditioned medium (UCM), set to 1. (e) P3 probe fluorescence measuremen t of selenide levels following mixture of selenite with 36 hr conditioned media from indicated cell lines. (f) Measurement of total selenium levels in selenite and vehicle/erastin treated NCIH838 and Hela cells. Total intracellular selenium was quantified following selenite supplementation for 2 hr to the 24 hr conditioned media with/without erastin. 3 µM erastin did not affect cell viability in 24 hr (Sup Fig 3k). (g) Total thiol quantification of conditioned media from CTRL, GSS KO, and GCLC KO SK-Hep1 cells treated with vehicle or 3 µM erastin with/without L-Cysteine. (h) P3 probe fluorescence measurement of selenide levels following mixture of selenite with 48 hr conditioned media from indicated cell lines. Measurement of selenide produced in control, GSS KO, and GCLC KO SK-Hep1 cells. (i) Quantification of total selenium in selenite and vehicle/erastin/L-Cysteine treated SK-Hep1 cells. Total intracellular selenium was measured following selenite supplementation with or without L-cysteine for 2 hr to the 24 hr conditioned media with/without erastin. Data are mean ± S.D. from biological replicates (nV=V3 for d-i) and were analyzed by two-tailed Student’s *t*-test. (d,e,g-i; \**P* < 0.05, \*\**P* < 0.01, \*\*\**P* < 0.001, *n.s.*, not significant).

### Selenide donates its electrons to ubiquinone

To begin to explore the selenoprotein-independent mechanism by which selenium is exerting its immediate lipid antioxidant / cytoprotective properties, we characterized the metabolome of cells treated with 6 µM selenite. Numerous metabolic pathways such as arginine and proline metabolism and aminoacyl-tRNA biosynthesis upon selenium treatment, were found to be significantly altered (Sup Fig 4a,4b, Sup Table S2). Most lipid species including PUFA-containing phospholipids and precursor polar metabolites for lipid synthesis were not significantly altered by 2 hr or 24 hr selenium treatment (Sup Fig 5a-5i); similarly, neither saturated lipids nor unsaturated lipids, as groups, were significantly altered due to selenium (Sup Fig 5f,5g). These results rule out lipid profile alterations as contributing to the anti­ferroptotic effect of selenium. ROS formation or iron levels were not affected by selenite treatment, ruling out these mechanisms also (Sup Fig 4f-4h). Selenide did not have any direct radical trapping activity that could explain its rapid antioxidant effect (Sup Fig 4i).

Examining the individual metabolites changed in our metabolic analyses, we observed that among the 165 metabolites measured, ubiquinol was among the few metabolites that were at least 2­fold-increased with significance (Fig 3a,3b, Sup Fig 4b). In particular, it was the top metabolite among the 15 metabolites in our analyses that are known to regulate ferroptosis, drawing our attention (Sup Fig 4c,4d). Ubiquinol is formed by the reduction of ubiquinone from multiple sources, and has emerged as an important antiferroptotic molecule due to its lipid peroxyl radical trapping activity ^16–18^. Thus, it was an attractive candidate to mediate the effects of selenium on ferroptosis independently of selenoprotein production.

**Figure 3.**
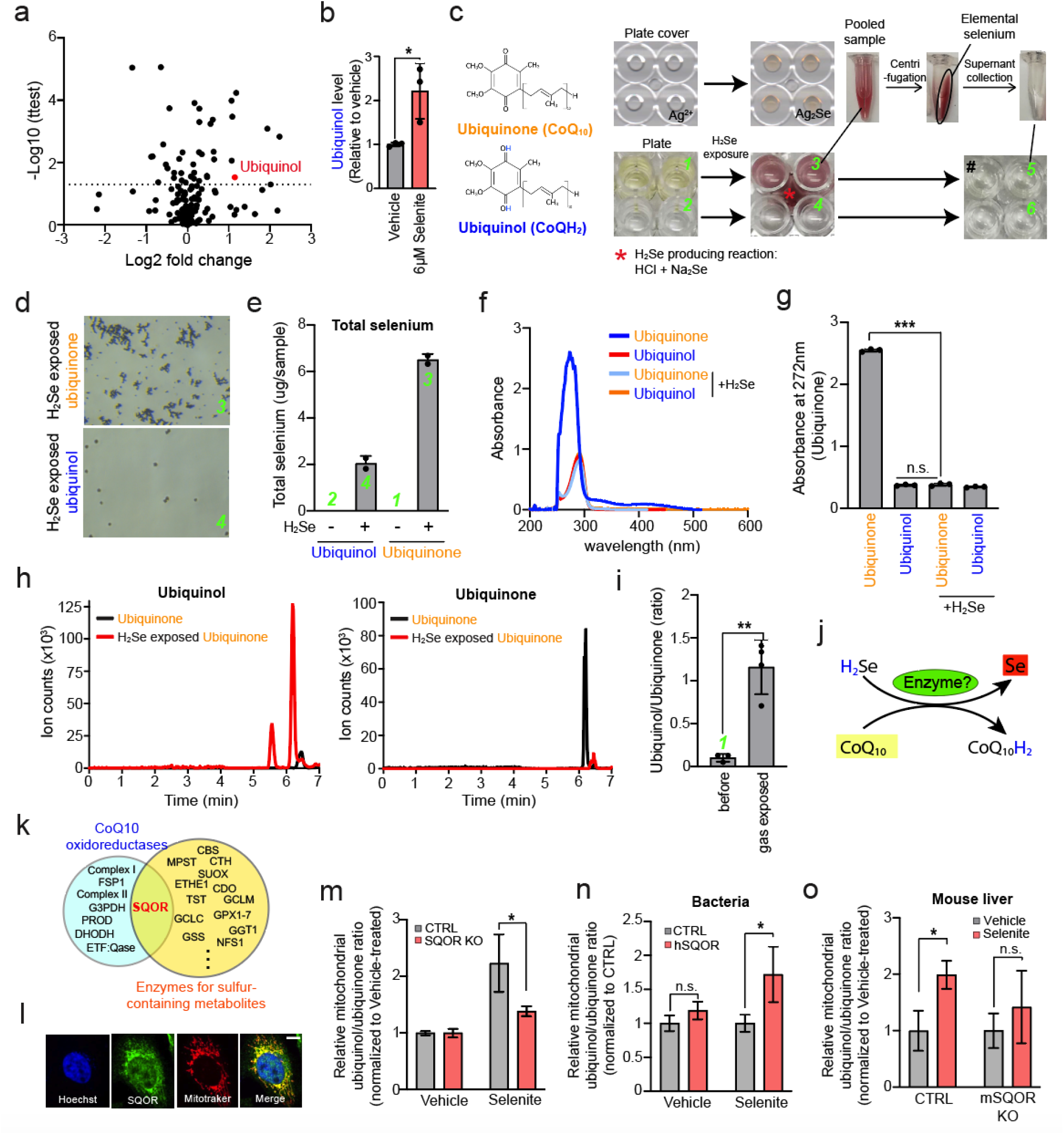
The selenium metabolite selenide reduces ubiquinone to ubiquinol via SQOR enzyme. (a) Scatter plot represents fold change of 165 metabolites and p-value (-log10 scale) of change upon 6 µM selenite treatment. (b) Ubiquinol/ubiquinone ratio of the selenite treated SK-Hep1 cells. (c) 96well plate cover spotted with silver nitrate embedded polyvinylpyrrolidone matrix to detect selenide (upper lef t images) and 96 well plate bottoms containing selenide gas producing solution (red asterisk*) and adjacent wells containing 200 µM ubiquinone and ubiquinol solutions (lower left images), before and after selenide gas exposure. The selenide gas exposed solutions were pooled in Eppendorf tubes and the elemental selenium was removed by centrifugation. The supernatant was placed into 96 well plate to compare the color of solution with ubiquinol (UQ_10_H_2_). The green numbers shown mark the sample s used to for various measurements as indicated in the following panels. (d) Brightfield images of selenide gas exposed 200 µM ubiquinone and ubiquinol solutions. 100X magnification. (e) ICP-MS measurement of total selenium in the ubiquinone and ubiquinol solution before and after gas exposure. (f) UV-vis spectrophotometer analysis of ubiquinone and ubiquinol before and after gas exposure. (g) Absorbance values of ubiquinone/nol at 272 nm, the peak wavelength which indicates ubiquinone quantity. (h) Ion counts of ubiquinol/none from LC-MS analysis in the ubiquinone solution before and after selenide gas exposure. (i) Ubiquinol/none ratio in ubiquinone solution before and after selenide gas exposure. (j) Predicted chemical reaction of hydrogen selenide and ubiquinone. (k) Venn diagram represents enzymes belonging to known CoQ10 oxidoreductases and sulfur processing enzymes^48, 49^. The full list is provided in Sup Fig 6e. (l) Immunocytochemistry of SQOR protein, mitochondrial marker (Mitotracker), and nucleus. Scale bar indicates 10 µm. (m) Relative ubiquinol/none (CoQ_10_) ratio in mitochondria from CTRL and SQOR KO SK-Hep1 cells treated with vehicle or 6µM selenite for 2 hr. (n) Relative ubiquinol/ubiquinone (CoQ_8_) ratio in control or human SQOR protein induced BL21 bacteria after treatment with 1µM selenite or vehicle for 2 hr. (o) Relative ubiquinol/none (CoQ_9_) ratio in mitochondria from wild type and SQOR KO mouse at 24 hr after 6mg/kg selenite intraperitoneal injection. Data are mean ± S.D. from biological replicates (*n* = 2 for e; *n* = 3 for a,b,g,i,m,n) and ± S.E.M. from biological replicates (n = 5 for o) and were analyzed by two-tailed Student’s *t*-test. (b,g,i,m,n,o; \**P* < 0.05,\*\**P* < 0.01,\*\*\**P* < 0.001, *n.s.*, not significant).

As selenide is a reducing agent ^19^, we wondered whether selenide could react with and reduce ubiquinone. To test this possibility, we produced selenide gas via a chemical reaction (Methods) in a central location of a 96-well plate and examined the reaction of selenide gas with neighboring wells containing solutions of ubiquinone (naturally yellow hue) and ubiquinol (clear) as candidate reactants (Fig 3c and Sup Fig 6a). Surprisingly, the wells containing ubiquinone, but not ubiquinol, turned red with colloid formation, suggesting the formation of elemental selenium (Se ^0^)^20^(Fig 3c and Sup Fig 6a). Microscopic images (Fig 3d and Sup Fig 6b), as well as quantification of selenium via ICP-MS (Fig 3e), verified that the ubiquinone-containing wells preferentially formed colloidal selenium, compared to the ubiquinol containing wells. Importantly, upon removal of colloidal selenium by centrifugation, it could be observed that the ubiquinone solution was no longer yellow but clear, raising the possibility that the ubiquinone had been converted to ubiquinol (Fig 3c). Ubiquinone has a distinct UV spectrum compared to ubiquinol, and our UV spectrum analyses validated a ubiquinone-to-ubiquinol conversion (Fig 3f,3g and Sup Fig 6c,6d). We further demonstrated the identity of the product in the well as ubiquinol via LC/MS (Fig 3h,3i). Overall, these data indicate a chemical reaction between selenide and ubiquinone, where electrons from selenide reduce ubiquinone, leading to the formation of electron-depleted Se and ubiquinol as products (fig 3j). While these experiments are not relevant to a cellular context, they demonstrate that selenium can donate electrons to ubiquinone, suggesting a means by which selenium could impact ferroptosis in a selenoprotein-independent manner.

### SQOR mediates selenide-driven ubiquinone reduction

We demonstrated that selenium could donate electrons to ubiquinol *in vitro*, but wondered whether this activity in the cell might be enzymatically catalyzed as endogenous selenide levels (Fig 3j) are unlikely to be as high used in the vitro reactivity assay. Metabolic enzymes are often known to be promiscuous^21–23^, processing substrates which share similar chemical structures. For example, FSP1 was recently found to process Vitamin K, owing to its similar chemical structure with ubiquinone ^21^. Noting the chemical similarity between hydrogen sulfide and hydrogen selenide that allowed us to utilize sulfide probes in the measurement of selenide (Fig 2b, Sup Fig 3a,3b), we wondered whether along similar lines, a sulfide processing enzyme could process selenide. Among enzymes known to utilize hydrogen sulfide, our attention was drawn to the enzyme sulfide quinone oxidoreductase (SQOR) as 1) it is an enzyme which reduces ubiquinone in mitochondria (Fig 3l)^24^, and 2) it derives the reducing electron from hydrogen sulfide (Fig 3k, Sup Fig 6e). Thus, if SQOR could similarly utilize selenide as substrate, it could be mediating the selenium induced production of ubiquinol, and thus be involved in the ‘noncanonical’ function of selenium.

To determine whether indeed SQOR is catalyzing ubiquinol production by selenium, we examined ubiquinol production in SQOR KO cells, and found that selenite-induced mitochondria ubiquinol (CoQ_10_H_2_) production was abrogated (Fig 3m, Sup Fig 6f). We next expressed human SQOR in BL21 bacteria and found that its expression allowed selenite-induced ubiquinol (CoQ _8_H_2_) production to occur in bacteria (Fig 3n, Sup Fig 6h,6i). Finally, we set out to examine the relevance of this mechanism *in vivo* . As selenite delivered orally has limited bioavailability ^25, 26^, we delivered selenite via intraperitoneal injection. This was able to significantly induce ubiquinol (CoQ _9_H_2_) formation in mouse liver (Fig 3o, Sup Fig 6g). Importantly, this effect was ablated in *mSQOR* knockout mice. Collectively, our results demonstrate that SQOR mediates selenium’s propensity to produce of ubiquinol in cells.

### SQOR protects against ferroptosis

As SQOR mediates the formation of the peroxyl radical trapping ubiquinol, it may be an important negative regulator of ferroptosis. Therefore, we examined whether modulation of SQOR can impact ferroptosis sensitivity in 10% serum media (with basal levels of selenium, and without additional selenite supplementation). Indeed, the disruption of SQOR sensitized cells to lipid peroxidation and ferroptosis induced by GPX4 inhibitors (Fig 4a-4d, Sup Fig 7b), while its overexpression was protective against ferroptosis (Sup Fig 7a). We noted that the effects of SQOR OE or KO were only partial in protecting or sensitizing against ferroptosis, likely reflecting how SQOR effects would be limited to the mitochondria while RSL3, as an inhibitor of GPX4 species across the cell, causes lipid peroxidation both inside and outside of mitochondria^16, 27, 28^.

**Figure 4.**
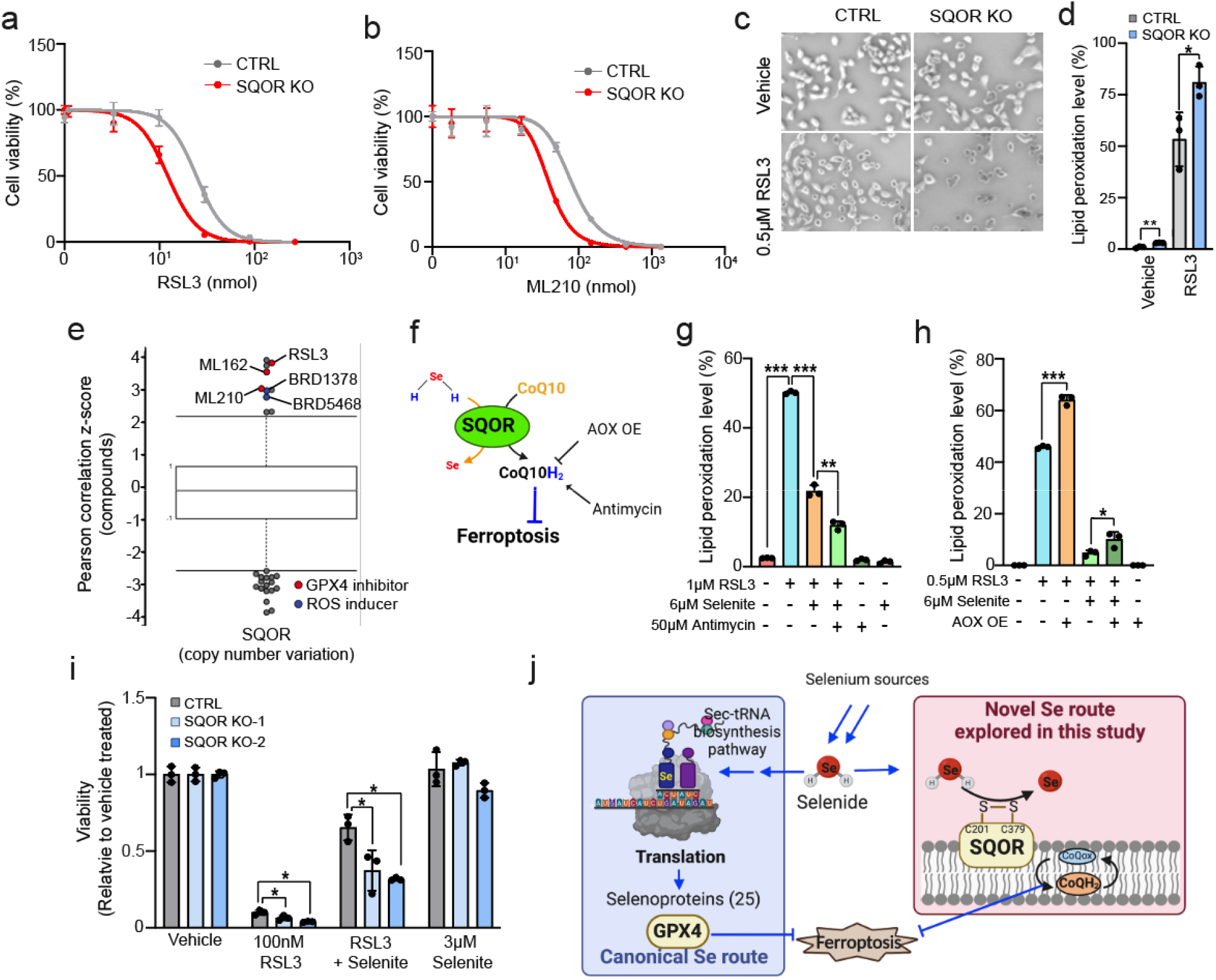
SQOR protects against ferroptosis. (a, b) RSL3 (a) and ML210 (b) dose response curve for CTR L and SQOR KO SK-Hep1 cells. Cell viability was measured at 24 hr after treatment with the GPX4 inhibitors. (c) Brightfield image of CTRL and SQOR KO SK-Hep1 cells treated with vehicle or 0.5 µM RSL3 for 2 hr. (d) Lipid peroxidation levels in CTRL and SQOR KO SK-Hep1 cells treated with vehicle or 0.5 µM RSL3 for 2 hr. (e) High SQOR copy number correlates with resistance to GPX4 inhibitors (RSL3, ML162, and ML210) and ROS producing drugs (BRD1378 and BRD5468) in cancer cells. Cell line drug sensitivity data were mined with Cancer Therapeutics Portal v2^50^. Each dot represents different drugs and the plotted values are z-scores of Pearson’s correlation coefficients of drug sensitivity with SQOR cop number across the 834 different cell lines. (f) Schematic diagram of approaches to modulate ubiquinol levels. Alternative oxidase (AOX) expression lowers, and complex III inhibitor Antimycin increases, the levels of ubiquinol formed by the SQOR-selenide axis, respectively. (g) Lipid peroxidation levels in vehicle or/and selenite treated SK-Hep1 cells with/without Antimycin and with/without RSL3 treatment, concomitantly for 2 hr. (h) Lipid peroxidation levels SK-Hep1 cells with/without induced AOX overexpression, treated with/without RSL3 for 2 hr. (i) Viability of CTRL and SQOR KO SK-Hep1 cell s treated with RSL3 and/or selenite for 24 hr. (j) A schematic diagram of how SQOR mediated ubiquinol production is the mechanism explaining how selenium suppresses ferroptosis in a selenocysteine - independent manner. Data are meanA±AS.D. from biological replicates (*n* = 3 for d,g,h,i) and were analyzed by two-tailed Student’s *t*-test. (d,g,h,i; \**P* < 0.05,\*\**P* < 0.01,\*\*\**P* < 0.001).

As SQOR could be generating ubiquinol from hydrogen sulfide, we tested and found SQOR loss still sensitizes to RSL3 toxicity when sulfide production is inhibited, indicating the relevance of selenium in these contexts (Sup Fig 7c-7f). In addition, RSL3 toxicity in SQOR KO cells was prevented by ferrostatin-1, a lipophilic antioxidant, and Mitotempo, a mitochondrial ROS scavenger (Sup Fig 7g,7h), and mitochondrial localized peroxidized lipids were relatively higher in RSL3 treated SQOR KO cell than control (Sup Fig 7i,7j), further supporting that SQOR negatively regulates mitochondrial ferroptosis. Unbiased drug sensitivity data analysis across a large panel of cancer lines demonstrates that cancer lines with high SQOR copy number are more resistant to GPX4 inhibitors (RSL3, ML162, and ML210) as well as ROS inducing compounds (BRD1378, BRD5468) (Fig 4e), suggesting a role for SQOR as a guardian against ferroptosis across cancer cells.

If the noncanonical, immediate selenium lipid antioxidant function involves mitochondrial ubiquinol production, then mechanisms which increase or remove mitochondrial ubiquinol should either potentiate or negate selenium’s effects (Fig 4f). Indeed, antimycin, a complex III inhibitor which thus impairs the shunting of ubiquinol into electron transport, enhanced the immediate suppression of lipid peroxidation by selenite at 2 hr (Fig 4g, Sup Fig 8a). The protective effect of antimycin against ferroptosis is in agreement with previous reports of antimycin protecting cells against reactive oxygen species-mediated death from xCT impairment or glutamate^28, 29^.

Conversely, overexpression of alternative oxidase (AOX), a yeast enzyme that can oxidize ubiquinol when ectopically expressed in mammalian cells ^30–32^, largely negated the noncanonical selenium antioxidant mechanism, indicating that ubiquinol is involved (Fig 4h, Sup Fig 8b-8d).

Based on these findings, we set to determine whether SQOR was responsible for the selenoprotein independent, anti-ferroptotic effects of selenium that we observed earlier. SQOR KO/OE by itself did not impact the expression of key anti-ferroptosis proteins such as FSP1, GPX4, and DHODH, and other CoQ10 Oxidoreductases (Sup Fig 8e,8f) and mitochondrial mass (Sup Fig 8g,8h), ruling these out as possible contributory factors. Importantly, the anti-ferroptotic effect of selenite was significantly diminished in SQOR KO cells (Fig 4i). Notably, there was some protection by selenium even under SQOR KO, likely representing the effects of selenoprotein induction which is in play in these 24h. DHODH is another mitochondrial CoQ10 reductase and thus a negative regulator of ferroptosis ^16^, and indeed we found DHODH loss sensitizes cells to RSL3 (Sup Fig 8i-8k). However, selenite could still rescue under these conditions (Sup Fig 8i-8k), further supporting that it is not working through DHODH. Thus, selenium potently suppresses ferroptosis via SQOR-catalyzed ubiquinone reduction. Taken together, these data demonstrate that the cytoprotective role of selenium extends beyond canonical selenoprotein production and also involves the production of ubiquinol via SQOR, which gives selenium a direct and rapid lipid antioxidant effect (Fig 4j).

## Discussion

The canonical role of selenium is to make selenoproteins, and because glutathione peroxidases and thioredoxin reductases are selenoproteins, selenium is often looked at as an antioxidant and cytoprotectant^1^. Here, we demonstrate selenium has a previously unknown role via a novel biological mechanism that is completely unrelated to selenoproteins. Selenide formed during the selenocysteine synthesis metabolic pathway deposits electrons to ubiquinone, generating ubiquinol in the mitochondria to suppress lipid peroxidation and prevent ferroptosis. The selenide mediated ubiquinol formation can be catalyzed by a highly conserved enzyme (SQOR), suggests this to be a fundamental aspect of selenium that may be conserved throughout biology.

Future efforts should explore how to leverage this mechanism towards therapeutic means. Lipid peroxidation-induced death is relevant to cellular demise in a variety of contexts including stroke, myocardiopathy, and excitotoxicity^5–7^. For example, glutamate excitotoxicity involves mitochondrial ROS and subsequent Bid translocation ^33, 34^. We observed that selenide prevents glutamate toxicity as well as downstream Bid translocation (Sup Fig 9), further implicating it is a protector against diverse ferroptotic contexts. On the other hand, we note that selenide *per se*, due to its volatile nature and toxicity at high or prolonged doses, is unlikely to be a practical therapeutic strategy. An attractive option for maximizing selenium distribution while minimizing toxicity is the use of cell permeable peptides containing selenocysteine residues, which have demonstrated protective properties in *in vivo* contexts such as stroke^5^. Maximizing selenium delivery, as well as SQOR mediated intracellular conversion to ubiquinol, may provide an attractive alternative anti-ferroptotic strategy compared to, for example, administration of CoQ10. While CoQ10 administration strategies have been shown promise in disease contexts ^35, 36^, they have not translated to clinical success, which may be in part due to limited bioavailability/delivery of CoQ10 supplementation in the human body^25, 26^.

In conclusion, our finding expands the view of selenium in cytoprotection, adding a direct and rapid mechanism, and at the same time identifying a novel selenium processing enzyme. In addition to the canonical role of selenium in allowing translation of selenocysteine-containing selenoproteins, and in triggering an antioxidant selenoprotein transcription program, we now show that selenium has an immediate and highly meaningful effect via enzyme-catalyzed production of ubiquinol in the mitochondria. Our findings reveal selenium to be an element that is emerging as having a more complex biology than previously appreciated.

## Methods

Information for all materials and oligos used in this paper is listed in Supplementary Table S1.

### Cell lines and cell culture

All cell lines were cultured in a humidified incubator at 37°C under 5% CO_2_ and atmospheric oxygen. SK­Hep1, Hela, NCIH838, SNU449, A498, DU145, LN229 and HT22 cells were cultured in Dulbecco’s modified Eagle’s medium (DMEM) with 10% (volume/volume; v/v) heat-inactivated FBS and 1% (v/v) penicillin/streptomycin. All cells were negative for mycoplasma test and verified by short tandem repeat (STR).

### Molecular cloning of expression constructs

SQOR was PCR amplified from cDNA produced from SK-Hep1 cells, using primers containing BamH1 and NOT1 sites for forward and reverse primers, respectively. The verified amplicon sequences corresponded exactly to the published consensus sequence for SQOR (CDS ID: CCDS10127.1, Size of amplicon: 1353bp). The amplicon was cloned into the pLV-EF1α-IRES-Blast vector. A catalytic mutant form of SQOR (C379A) was constructed by site-directed mutagenesis by Genewiz, and the sequence encoding cysteine (TGT) for 379^th^ amino acid was confirmed to be changed to the encoding sequence for alanine (GCC). All primer information is provided in Sup Table S1.

### Lentivirus production and CRISPR/Cas9-mediated genome editing in cell lines

Lentivirus was produced by transfecting pLentiCRISPR v2 ^37^ containing guide sequences targeting SQOR, PSTK, GSS, GCLC, CBS, CSE, or MPST genes, pLV-EF1α-IRES-Blast vector containing SQOR gene sequence, or 48 hr. pCW57.1_yeastAOX ^32^ with the Delta-Vpr packaging plasmids and the VSV-G envelope plasmid (DNA concentration ratio, 1.2: 1: 0.3) into HEK293T cells using X-tremeGENE 9 Transfection Reagent (Roche). Lentivirus-containing media was harvested at 24 hr after fresh media change. The supernatant containing the virus was directly used without concentration after one cycle of freeze and thaw to kill off any residual HEK293T cells in the supernatant. The virus was transduced into the target cell lines with 10 µg/ml polybrene for 24 hr, and then the media was replaced with media with puromycin (typically at 1 µg/ml) or blasticidin (typically, 10 µg/ml) for cells infected with the virus containing pLentiCRISPR v2/pCW57.1_AOX or with the virus containing pLV-EF1α-IRES-Blast vector, respectively, to select for transduced cells. For the experiment involving the expression of AOX, single-cell clones were treated with 500 nM doxycycline (Sigma) for 48hr to induce its expression. All guide sequences information is provided in Supplementary Table S1.

### Western blot analysis

Immunoblotting to confirm protein expression was performed as previously described ^38^. In brief, cells were lysed using RIPA lysis buffer (Boston bioproducts) supplemented with protease inhibitor (Roche), and the protein concentration was determined using a Bradford Assay (Biorad) with a Beckman Coulter DTX880 Multimode plate reader. For the bacteria, the pellets were resuspended in 50 mM HEPES with protease inhibitor and lysed by sonication (1 sec on - 1 sec off for 30 sec, rest for 30 sec, 6 rounds total). Identical amounts of protein (typically, 15-30 µg) for each sample were denatured, boiled in 6X Laemmli buffer (Boston bioproducts) at 90°C for 5 min, and loaded on a polyacrylamide gel. The standard immunoblotting was performed, and ECL, Pico, or diluted Femto (Pierce) substrate was used for detection. The antibody information is provided in Sup Table S1.

### Cell viability assay

Cell viability was performed with Hoechst /PI (propidium iodide) double staining or CellTiter-Glo Luminescent Assay.

### Hoechst/PI double staining

The drugs were treated at 24 hr after cell seeding (1500 cells/well in 96 well plate). At the indicated time point, media was replaced with 100 µl phenol-read free media containing 1 µg/ml PI (Sigma Aldrich) and 2 µM Hoechst (Thermo Fisher Scientific). After 20 min incubation at 37°C, fluorescence images (blue channel for Hoechst and red channel for PI) were obtained, and the number of Hoechst stained and PI-stained cells were counted using a cell viability application of Celigo Imaging Cytometer. The number of hoechst stained cells and PI-stained cells represent total cell number and dead cell number, respectively. The total cell number (the number of hoechst-stained cells) subtracted by dead cell number (the number of PI-stained cells) was defined as viable cell number. The viable cell number of drug treated condition was normalized to the viable cell number of vehicle treated one (vehicle treated condition was set as 1). The viability data, Sup Fig 2d,2f, Fig 4a,4b, and Sup Fig 7a were obtained with Hoechst/PI double staining method.

### CellTiter-Glo Luminescent Assay

20 µl of CellTiter-Glo (CTG) luminescent solution (Promega) was added in each well after equilibration of the 96 well plate and CTG solution at room temperature for 20 min. Luminescence signal was captured using a Beckman Coulter DTX880 Multimode plate reader with a setting at 1.0 sec. The values were normalized to the values from vehicle-treated cells. Fig 1d,1i, Fig 4i, Sup Fig 2e,2g,2h, Sup Fig 3k, Sup Fig 7c-7h and Sup Fig 9a were obtained with CTG assay.

### Lipid peroxidation measurement

180,000 cells/well were seeded on 6-well plates and incubated in a CO_2_ incubator One day before the assay. Cells were treated with RSL3 and/or selenite in the fresh media and incubated in the CO_2_ incubator. After 2 hr incubation, single cells were collected by trypsinization and resuspended in 150 µl EBSS (Thermo Fisher) containing 5 µM C11-BODIPY 581/591 (Invitrogen), a lipid-soluble fluorescent indicator of lipid oxidation, or MitoPerOx (Abcam), a fluorescent mitochondria-targeted lipid peroxidation probe. After 20 min incubation in the CO_2_ incubator, lipid peroxidation was quantified using the flow cytometer BioradZE5 with FL03 laser. 10,000 single cells were used for the analysis per sample. The gating strategy is presented in Sup Fig 1h.

### Usage of selenide/selenite solution

Depending on the purpose and time, or feasibility of the assays, sodium selenite (selenite) solution, sodium selenide (selenide) solution, or hydrogen selenide (selenide gas) was used. In this study, selenide gas was used for confirming the ubiquinone reduction to ubiquinol by hydrogen selenide in Fig 3c-3i. The detailed method of how to produce selenide gas is described in a separate section, “production of selenide gas”. Selenide gas is produced rapidly by mixing the sodium selenide in an acidic (3.7%HCl) solution, and diffuses to surrounding wells in a circular pattern, and turns red according to whether elemental, colloidal selenium is formed. This allows us to ascertain and compare/contrast chemical reactivity of the solutes dissolved in the various wells upon exposure to the gas. This method produces a high but short-lived burst of selenide gas production, and thus selenite or selenide solution was used instead for the cell-based assays such as viability assay, BODIPY dye lipid peroxidation quantification and metabolites analysis. In the water (rough pH is between 6.5 and 8.5), selenide is present in a form of selenide anion (HSe^-^)^39^, which is also not as stable as selenite in water. Thus, the selenide stock solution using selenide powder (light pink color) was freshly made for every experiment.

### Production of selenide gas

As sodium selenide powder (light pink color) is hygroscopic and easily oxidized (when it is oxidized, the color is changed to dark brown, gray, or black), the powder was aliquoted into glass insert loaded glass vial which is used for loading samples for mass spectrometry (MS) and tightly sealed with parafilm. When selenide powder is used, it is important to minimize the time for selenide exposure to the air. Hydrogen selenide gas was formed by mixing fresh sodium selenide around 50 mM concentration, into 3.7% hydrochloric acid solution (HCL+Na _2_Se→H_2_Se+2NaCl). Since hydrogen selenide gas is volatile, 100 µl of the mixture was placed into the center of 96 well plate right after mixing the sodium selenide with hydrochloride acid. Immediately following this, the plate cover containing silver-PVP spot was placed on the plate bottom to confirm the selenide gas production as described in the “hydrogen selenide gas detection” section, or the gas source was placed into the 96 well plate bottom containing wells filled with ubiquinone/nol solution to monitor the color changes of the ubiquinone/nol solutions by the selenide gas (Sup Fig 6a). All procedures were performed in the safety hood due to the toxic and volatile nature of selenide that was produced.

### Selenide detection

Three methods, the metal-embedded matrix selenide detection method we previously conceived based on the chemical similarity between sulfur and selenium ^9^, the selenide testing strip-based method, and a P3 fluorescent-based probe widely used for sulfide detection ^11^ but unexplored for selenide detection were applied to detect and quantify selenide formation. The P3 fluorescent-based probe was used for cell based assay as it showed better sensitivity.

### Metal-embedded matrix selenide detection method

As described previously, silver embedded matrix can be used for selenide gas detection ^9^, and utilized for Fig 3c. Briefly, 5% polyvinylpyrrolidone (PVP) solution (wt/vol) was mixed with 1M silver nitrate solution at a ratio of 9:1, yielding final concentrations of 100 mM of silver nitrate, and 4.5% PVP. The silver-embedded PVP solution was mixed well, and then 15 µl of silver–PVP mixture was dropped on the inside of the cover of a transparent 96-well plate. To solidify the matrix solution, the cover was dried at room temperature for at least 1 hr after covering with aluminum foil which minimizes the silver oxidation. This plate cover having the ‘silver-matrix spots’ was placed above the 96 well plate bottom immediately after hydrogen selenide gas producing mixture was added into the center of the plate. The spots produced brownish silver selenide precipitate (H_2_Se+2AgNO_3_→Ag_2_Se+2HNO_3_; Fig 2d), which represents successful production of hydrogen selenide gas. After 1 min exposure, plate cover was displaced from the source, stayed in the safety hood to let any residual selenide gas removed for 10 min, and then scanned.

### Selenide testing strip-based method

The lead acetate-embedded testing strip (Industrial Test System) was dipped into freshly prepared selenide standard solution for 1 sec and brown color was visualized in a dose-dependent manner as selenide reacts with lead to form brown lead selenide (H _2_Se+ Pb(C_2_H_3_O_2_)_2_→Pb_2_Se+2(C_2_H_3_O_2_); Fig 2b). To test the selenide production capacity of thiol and non-thiol­containing metabolites, the detection paper was dipped into the selenite solution right after mixing with the metabolite solutions for 1 sec and the paper was left to sit for 1 min and the picture was taken.

### P3 fluorescent-based probe

We confirmed that Sulfide P3 fluorescent probe could be used as a selenide indicator by measuring freshly prepared selenide standard solutions, which increased fluorescent signal in a dose-dependent manner (Sup Fig 3b). To confirm the dose curve of P3 fluorescence with selenide, 5 µl of 20 mM P3 probe dissolved in DMSO (final working concentration of P3; 1 mM) was diluted in 85 µl water and then mixed with 10 µl freshly prepared selenide standard and incubated for 10 min. Next, the fluorescent signal was measured with Glowmax plate reader (Emission: 500-550 nm, Excitation: UV365 nm). For testing selenide production capacity of thiol and non-thiol containing metabolites, each metabolite was mixed with 500 µM selenite and 1 mM P3 probe in total volume of 50 µl. For testing cellular selenide production capacity, 50,000 cells/well were plated in a 96 well plate. Next day, growth media was replaced with phenol red free media with or without 3 µM erastin. After 36 hr of media conditioning, 75 µl of the phenol red free growth media was mixed with 10 µl of 1 mM P3 and selenite solution (final effective concentration of P3 and selenite; 117 µM). After 10 min incubation in 37°C incubator, the fluorescent values were obtained. All values were subtracted by that of blank to remove noise from media or water.

### Intracellular iron measurement

180,000 cells/well were plated on 6-well plates one day before the assay. Cells were treated with selenite or 2,2-bipyridyl (Bpy) (Sigma) in fresh media and incubated in the CO_2_ incubator. Bpy was used as a chelator of iron. After 2 hr incubation, cells were collected by trypsinization and resuspended in 150 µl EBSS (Thermo Fisher) containing 1 µM FerroOrange (Dojindo), a fluorescent indicator of iron. After 20 min incubation in the CO_2_ incubator, iron level was quantified using the flow cytometer BioradZE5. 10,000 single cells were used for the analysis per sample. The gating strategy is presented in Sup Fig 1h.

### Superoxide measurement *in vitro*

To measure radical trapping activity, superoxide radical was generated by mixing 50 µM H_2_O_2_ with 1 µM iron. First, 50 µl of water or 2XH _2_O_2_ was added to a 96 well plate and 50 µl of 20 µM superoxide radical indicator MitoSOX (ThermoFisher) probe. Then, 1 µl of iron solution to generate the fenton reaction creating superoxide, and 1 µl of selenide or superoxide dismutase (SOD) enzyme as a positive control was added for testing the radical trapping activity. After 10 min incubation at 37°C, the fluorescent signal was measured with 580-640 nm emission and 530 nm excitation filter (GloMax plate reader).

### ROS measurement

180,000 cells/well were plated on 6-well plates. Next day, SK-Hep1 cells were treated with vehicle or 6 µM selenite (Santa Cruz) in fresh media and incubated in the CO_2_ incubator. HT29 cells were treated with 5 mM Glutamate and/or 3 µM selenide (Santa Cruz). After 2 hr and 24 hr incubation for SK-Hep1 and 12hr for HT29 cells, cells were collected by trypsinization and resuspended in 150 µl EBSS (Thermo Fisher) containing 10 µM DCFDA (Abcam), a fluorescent indicator of ROS. After 30 min incubation in the CO_2_ incubator, the ROS level was quantified using the flow cytometer BioradZE5. 10,000 single cells passing through the gating strategy provided in Sup Fig 1h were used for the analysis per sample.

### Mitochondria content measurement

180,000 control and SQOR OE/KO cells/well were plated on 6-well plates and incubated in the CO_2_ incubator. The next day, one extra well was replaced with 1% FBS media as a serum-starved condition to induce mitochondrial mass increase, which was used as a positive control. After 36 hr incubation in the CO_2_ incubator, cells were stained with 200 nM Mitotraker Deep Red FM (Cell Signaling Technology) in the media for 30 min, and then cells were collected with trypsinization. The collected cells were resuspended with phenol red-free media, and then total mitochondrial mass was quantified using the flow cytometer BioradZE5. The gating strategy for 10,000 single cells read is indicated in Sup Fig 1h.

### Mitochondria membrane potential measurement

180,000 HT29 cells/well were plated on 6-well plates. Next day, treated with 5 mM Glutamate and/or 3 µM selenide (Santa Cruz). After 12 hr, cells were collected by trypsinization and the cell pellet was resuspended within 500µl of 400nM TMRE (Abcam), a positively-charged dye that readily accumulates in mitochondria because of their relative negative charge with accumulation correlating to the degree of membrane potential, and 10% FBS (v/v) contained phenol red-free DMEM (Thermo Fisher). After 25 min incubation in the CO_2_ incubator, the TMRE intensity of single cells was quantified using the flow cytometer Biorad ZE5F with FL04 laser. During the live-cell analysis, the samples were kept at 37°C. 10,000 single cells were used for the analysis per sample. The gating strategy is presented in Supplementary Figure .

### Conditioned media total thiol quantification

Total thiols in the conditioned media was assessed by Ellman’s test ^38^. 50,000 cells/well for Fig 2d,2e or 10,000 cells/well for Sup Fig 3h were plated in 96 well plate. The next day, the media was replaced with 100μl fresh phenol red free media with or without vehicle or 3 μM erastin. After 30 hr conditioning for Sup Fig 3h or 24 hr conditioning for Fig 2d,2e, 50 μl of conditioned media was directly mixed with 25 μl of 10 mM DTNB (5,5-dithiobis-(2-nitrobenzoic acid)) dissolved in DMSO. Colorimetric changes was measured 3 min after mixing the conditioned media with DTNB solution at 450 nm of absorbance (DTX880, Beckman Coulter, Indianapolis, IN, USA). The value from DTNB only was substracted as noise, and all values from the mixture of media and DTNB were normalized to that of unconditioned phenol free media. The leftover cells which were used for media conditioning in 96 well plate were used for the CellTier-Glo Luminescent Assay.

### Subcellular organelle fractionation

Subcellular organelle fractionation was carried out with Cell Fractionation Kit (Abcam). Briefly, 2 million cells/plate were plated in 100pi dish plates. The next day, the media was replaced with fresh growth media containing vehicle or selenite. After 2 hr and 24 hr of incubation, cells were collected by trypsinization. The cell pellets were reconstituted with 25 0μl of Buffer A and 250 μl of Buffer B containing Detergent I was mixed in by pipetting. The mixture was incubated on a rotator at room temperature. After 7 min incubation, the mixture was centrifuged at 5000g at 4°C for 1 min, and the supernatant was collected and centrifuged again at 10,000g for 1 min, and the pellet was kept on ice. The supernatant after two rounds of centrifugation was stored (cytoplasmic fraction). The cytoplasm-depleted pellets were reconstituted with 250 μl of buffer A, buffer C containing Detergent II was mixed into the sample tube, and then the mixture was incubated on a rotator at room temperature. After 10 min incubation, the mixture was centrifuged at 5000g at 4°C for 1 min, and the supernatant was collected (mitochondria fraction) and centrifuged again at 10,000g for 1 min, and the pellet (nucleus) was reconstituted with 250 μl of buffer A. All fractions were mixed with the same volume of 2XRIPA buffer and the samples were used for SDS-PAGE.

### Quantification of mRNA of selenoproteins

SK-Hep1 and U251 cells (400,000 and 800,000 cells/well for 24 hr and 2 hr, respectively) were seeded on 60pi cell plates. The next day, 6 μM or 3 μM selenite was treated, and cells were collected after 2 hr or 24 hr treatment of selenite. The SK-Hep1 and U251 cell pellets (biological triplicates each condition) were re-suspended in TRIzol reagent (Invitrogen), briefly vortexed, and chloroform (Sigma-Aldrich) was added. Following centrifugation at 12,000g at 4°C for 10 min which allowed for phase separation, the transparent supernatant was collected and mixed with isopropanol (Sigma-Aldrich). Further centrifugation in the same condition enriched the RNA pellets, and the pellets were washed three times with 70% EtOH and eluted in Nuclease-free distilled water (Invitrogen). From the RNA samples, cDNA was synthesized by using SuperScript First-Strand Synthesis kit (Invitrogen) according to manufacturer’s instructions. The Real Time-qPCR gene expression quantifications were performed in 96-well plates (Applied Biosystems) using the StepOnePlus 96-well Real-Time PCR System (Applied Biosystems). Reactions with cDNA sample were carried out in a total volume of 20 μl, comprising 10 μl 2XSYBR Green Master Mix (Applied Biosystems) and 0.25 μM of each primer. The primer information is provided in Sup Table S1.

### Immunocytochemistry

Cells were seeded on Mattak 8-well chambered cell culture-treated slides (Mattek). 24 hours later cells were treated with vehicle, 5 mM L-Glutamate, and/or 3 μM selenide. After 12 hr of incubation, a 200 nM Mitotracker probe (Cell signaling) was added to all cell treatment conditions and incubated at 37°C for 30 min. The cells were rinsed once with PBS and fixed with 4% paraformaldehyde in PBS for 12 min at room temperature. The cells were then rinsed three times with PBS and were permeabilized with 0.1% TritonX-100(v/v) in PBS for 2 min at room temperature. Cells were washed three times and blocked in 1% BSA(v/v) in PBS for 1 hour at room temperature. The cells were incubated in the primary antibody at 4°C overnight, rinsed three times with PBS, and then incubated with the secondary antibody for 45 min at room temperature in the dark. Cells were washed three times with PBS and mounted on the slides using ProLong Gold Antifade mountant containing DAPI (Thermo Fisher). Images were acquired on the Nikon Eclipse Ti2 confocal microscope. Images were analyzed using ImageJ (1.53q).

### Measurement of metabolites with mass spectrometry

#### 1. Measurement of total selenium with ICP-MS

200 μM ubiquinone and ubiquinol solutions before/after selenide gas exposure as described in the ‘Production of selenide gas’ section were collected into tubes without centrifugation to compare total selenium in gas exposed ubiquinone and ubiquinol solution (Fig 3e). For measuring selenium uptake capacity (Fig 2h,2i), the previously established method which we conceived was applied ^38^. Briefly, 5 million cells/plate were plated in three 150 pi dish plates (biological replicates for each plate). The next day media was replaced with fresh media with or without 3 μM erastin. One day after media conditioning, selenite (final concentration; 6 μM) was added onto each plate containing the conditioned media with or without L-cysteine (final concentration; 50 μM) and cells were harvested by trypsinization and washed with cold PBS three times after 2 hr incubation. The cell pellets were weighed for normalization.

The samples were further processed as described previously ^9, 38^. Briefly, all samples were dried and then treated with 500μl of H_2_O_2_/HNO_3_ solution. After 12 hr incubation at RT with periodic venting, the samples were sonicated for one hour at 35kHz 40 °C with periodic venting, and then the samples in the e-tube were transferred to a glass digestion tube. Additional 0.5 ml DI rinses for three times in the same e-tube and the DI were also collected in the same digestion tube (total of 2 ml in the digestion tube). The tubes were sealed and boiled at 140°C. After 2 hr, the samples were cooled, and diluted with 10 ml with DI water for analysis.

All analyses were performed with an Agilent 7500A ICP-MS system fitted with a standard concentric nebulizer, a torch shield, a Peltier-cooled double-pass Scott-type spray chamber, and standard Ni interface cones. To transfer 10g l–1 yttrium as an internal standard, the peristaltic pump was used, and masses at 77–78, 82, 83 and 89 were obtained using a spectrum analysis (20 points per mass), with 3 repetitions. Using standard environmental correction for bromine (82=82–83), interference correction was performed on the mass at 82AMU. To minimize carry-over between analytical sample, the probe of the sample introduction system was washed with 10% nitric acid for 1 min. Calibration curves were obtained using standard selenium (Ultra scientific) solution and quality-control samples using a multielement standard (Environmental Express). Additionally, mock samples were processed for each calibration and subjected to the sample preparation for insuring adequate recovery. The recoveries were 100±10% for all cases (data not shown). The data with calibration solutions was analyzed without background subtraction using the 89 internal standard peak. In all cases, the linear regression afforded an r ≥0.996.

#### 2. Measurement of polar metabolites with LC/MS

##### MS sample prep

For polar metabolite extraction, 700,000 SK-Hep1 cells/wells were seeded on 6 well plates in 1.3 ml growth media. The cell seeding number and the volume of media was determined to make a similar cell density and media environment for the condition used for ubiquinone/nol measurement in 150pi dish as cell density and prior media conditioning may affect the efficiency of selenite to selenide reduction. The next day, the media was replaced with the 1.3 ml fresh media containing vehicle or 6 µM selenite. After 2 hr 30 min, the media was removed, and the cells were washed with 2 ml of cold PBS for two times. Then, metabolites were extracted by adding total 1 ml of polar metabolite extraction buffer (80% methanol: 20% water solution) at dry ice temperature. For more detail, 500 μl of the extraction buffer was added on the cell plate and then incubated in deep freezer for 10 min and then the cell plate was placed on the dry ice. The cells were detached by scrapping on the dry ice and then collected into prechilled e-tubes. The residual cells and metabolites were collected into the same e-tubes with additional 500μl of extraction buffer. The e-tubes containing the cells/metabolites were vortexed for 10 min in the cold room. Then the samples were centrifuged at 13,000rpm for 10 min to collect supernatant. The samples were dried down in a Refrigerated CentriVap Benchtop Vacuum Concentrator connected to a CentriVap-105 Cold Trap (Labconco). Immediately prior to running the samples, the metabolite pellets were resuspended in HPLC grade water (Thermo Fisher Scientific), vortexed for 10 min at 4°C, and then centrifuged at 16,000g for 10 min at 4°C. Metabolite lysates were subsequently loaded into LC-MS vials.

##### LC/MS analysis

A QExactive benchtop orbitrap mass spectrometer equipped with an Ion Max source and a HESI II probe coupled to a Dionex UltiMate 3000 HPLC system (Thermo Fisher Scientific) was used for all metabolite profiling experiments. The mass spectrometer underwent calibration on a weekly basis. 2μl of the samples were injected onto a SeQuant ZIC-pHILIC 5 μm 150x2.1 mm column after going through a 2.1x20 mm guard column (Millipore Sigma). The column oven was maintained at 25°C, and the autosampler was maintained at 4°C. Buffer A was comprised of 20 mM ammonium carbonate, 0.1% ammonium hydroxide and Buffer B was comprised of 100% acetonitrile. The chromatographic gradient was operated at a flow rate of 0.150 ml/min as follows: 0-20 min: linear-gradient from 80-20% B; 20­ 20.5 min: linear-gradient from 20-80% B; 20.5- 28 min: hold at 80% B. The mass spectrometer was run with the spray voltage set to 3.0kV, the heated capillary at 275°C, and the HESI probe at 350°C, in full-scan and polarity switching mode. The sheath, the auxiliary, and the sweep gas flows were 40 units, 15 units, and 1 unit, respectively. The resolution was set at 70,000, the AGC targeted at 1x10 ^6^, and the maximum injection time was set at 20 msec. The levels of metabolites were quantified by integrating peaks using the TraceFinder 4.1 software (ThermoFisher). Metabolites were identified with a 5ppm mass tolerance based on the retention time as determined by chemical standards.

#### 3. Measurement of Ubiquinone (CoQ) and Ubiquinol (CoQH_2_) with LC/MS

##### Preparation of cell pellets

20∼30M cells were plated into 150pi dish plates with 20 ml of growth media The next day, the media was replaced with the 20 ml of fresh media containing vehicle or 6 µM selenite. After 2 hr 30 min, the media was removed, and the cells were washed with 10 ml cold PBS. And then, the cells from two cell plates were collected into a 50 ml tube by scraping the cells within 10 ml of cold PBS per plate. To collect residual cells on the plates, the plates were washed with an additional 10 ml cold PBS which is collected into the same 50 ml tube. Then, cells were pelleted by centrifugation at 1500 rpm for 3 min at 4°C, resuspended with 1 ml of cold PBS, and moved to 2 ml size e-tubes. The cells were pelleted again by centrifugation at 2500rpm for 5 min at 4°C and the PBS was suctioned. The cell pellets were stored at −80°C until the mitochondrial isolation was performed.

##### Preparation of bacterial pellets

For testing the human SQOR protein-mediated ubiquinol (CoQ_8_H_2_) production by selenium in bacteria, pET-30a+_hSQOR construct was transformed into BL21 bacteria. The experimental scheme is depicted in Sup Fig 6h. A starter culture was prepared by overnight growth of bacteria after bacteria inoculation at 37°C with vigorous shaking overnight. The next day, 1 ml of starter culture was inoculated in 50 ml of LB media with 50 µg/ml kanamycin. When the optical density value reached 0.7, protein expression was induced by adding 0.5 mM isopropyl β-Dthiogalactopyranoside (IPTG; Sigma Aldrich). After 20h incubation at 15°C, 3 ml of bacteria was moved to bacteria tubes and vehicle or selenite was added. The tubes were placed in 37 °C incubator with vigorous shaking. After 2 hr incubation, 1 ml of samples were placed to the Eppendorf tube and centrifuged at 4,200g for 10 min to obtain pellets which were used for ubiquinol/none measurement.

##### Preparation of mouse liver tissues

The experimental scheme is depicted in Sup Fig 6g. For testing SQOR-mediated ubiquinol (CoQ_9_H_2_) production by selenite supplementation, C57BL/6N-Sqor ^em1(IMPC)Wtsi^/WtsiH strain was obtained from (Wellcome Trust Sanger Institute). 6mg/kg selenite was intraperitoneally injected into 4 weeks old five WT and SQOR KO mice. After 24 hr, the mouse was sacrificed, the liver tissues were obtained and excised, and snap-frozen for mitochondrial isolation. The all mouse-related procedure was compliant with and were approved by the Institute Animal Care and Use Committee (IACUC) of University of Massachusetts Medical School.

##### Mitochondrial isolation

The cell pellets were re-suspended in 2 ml of mitochondrial isolation buffer (200 mM sucrose, 10 mM Tris HCl, 1 mM EGTA/Tris, protease inhibitor cocktail (Roche), pH7.4 (adjusted with 1M HEPES)), transferred into an ice-cold round-bottom polypropylene test tubes, and dounced with 13 strokes using a 3 ml syringe and 25-gauge needle. The frozen mouse liver tissues (around 30mg) were homogenized by 25 dounces in 2 ml of mitochondrial isolation buffer. Homogenates were transferred into prechilled 2 ml tubes and centrifuged at 600g for 10 min at 4°C. Supernatants were moved to prechilled tubes and centrifugation was repeated. Then, supernatants were collected to a new prechilled tube and centrifuged for pelleting at 7,000g for 10 min at 4 °C. Pellets were subsequently stored at −80°C until further analysis.

##### Metabolite extraction and LC/MS analysis

The mitochondrial and bacterial pellets were resuspended in 500μl of ethanol and vortexed for 10 min at 4°C. Next, 1 ml of hexane was supplemented, and the sample tubes were vortexed again for 10 min at 4°C. The sample tubes were centrifuged at 16,000g for 10 min at 4°C, forming two layers – hexane and ethanol for the top and bottom layers, respectively. The top (hexane) layer was collected into a new tube and dried down for LC-MS analysis. Right before LC-MS analysis, the dried metabolites were reconstituted in 50μl of ethanol:hexane (80:20) mix and loaded into a 4°C autosampler. 5 µl of samples were injected onto a Luna 3 µm PFP ^40^ 100 Å, LC Column 100 x 2 mm. The column oven temperature was 25°C. Buffer A (water with 0.1% formic acid) and Buffer B (acetonitrile with 0.1% formic acid) were used for the gradient elution of LC. The gradient elution condition was as follows: 0 to 3 min, hold at 30% Buffer A, 3 to 3.25 min, gradient to 2% Buffer A, 3.25 to 5 min, hold at 2% Buffer A, 5 to 6 min, gradient to 1% Buffer A, 6 to 8.75 min, hold at 1% Buffer A, 8.75 to 9 min, gradient to 30% Buffer A and 9 to 10 min, hold 30% Buffer A. The chromatographic gradient elution was run at 0.5 ml/min flow rate. The mass spectrometer was run in positive-ion mode, full scan, with the heated capillary at 275°C, the spray voltage set to 4.0kV, and the HESI probe at 350°C. The sheath, the auxiliary, and the sweep gas flows were 40, 15, and 1 units, respectively. MS data were obtained in a range of m/z = 500 –1000. The resolution was set at 140,000, and the AGC targeted at 3×10^6^, and the maximum injection time was set at 250msec. The ubiquinone/nol were quantified by integrating peaks using the TraceFinder 4.1 software (ThermoFisher). The ubiquinone/nol were identified with a 5ppm mass tolerance based on the retention time as determined by ubiquinone/nol standards.

#### 4. Lipidomics analysis

##### Sample preparation

400,000 cells/plate for 24 hr treatment of selenite/vehicle or 800,000 cells/plate for 2 hr treatment of selenite/vehicle cells were plated in 60pi cell plate. One day after cell seeding, the media was replaced with or without selenite. After 2 hr or 24 hr incubation, media was removed, and the cells were washed with 5 ml PBS once and cells were collected by scraping with 5 ml cold PBS. The cell suspension in PBS was pelleted by centrifugation at 1500rpm, 4°C for 3 min. The pellet was washed again with 500μl cold PBS, and the cell suspension was moved to e-tubes. The cell suspension was pelleted by centrifugation at 1500rpm, 4°C for 5 min and was stored at −80C for the lipid extraction. Before extraction, cell pellets were thawed on ice for 30 min and resuspended in 50μl PBS. Internal standards (5 µl SPLASH® LIPIDOMIX® per sample) dissolved in methanol were added directly to each suspension. Lipids were extracted by adding tert-butyl methl ether (1250 μl) and methanol (375 µl). The mixture was incubated on an orbital mixer for 1 hr (room temperature, 32rpm). To induce phase separation, H_2_O (315 µl) was added, and the mixture was incubated on an orbital mixer for 10 min (room temperature, 32rpm). Samples were centrifuged (room temperature, 10 min, 17,000g). Upper organic phase with collected and subsequently dried *in vacuo* (Eppendorf concentrator 5301, 1ppm).

##### Liquid Chromatography

Dried lipid extracts were reconstituted in chloroform/methanol (150μl, 2:1, v/v) and 15μl of each extract was transferred to HPLC vials containing glass inserts. Quality control samples were generated by mixing equal volumes of each lipid extract followed by aliquotation in 15μl aliquots. Aliquoted extracts were dried *in vacuo* (Eppendorf concentrator 5301, 1 ppm) and redissolved in 2­propanol (15μl) for injection. Lipids were separated by reversed phase liquid chromatography on a Vanquish Core (Thermo Fisher Scientific, Bremen, Germany) equipped with an Accucore C30 column (150 x 2.1 mm; 2.6 µm, 150 Å, Thermo Fisher Scientific, Bremen, Germany). Lipids were separated by gradient elution with solvent A (MeCN/H_2_O, 1:1, v/v) and B (i-PrOH/MeCN/H_2_O, 85:10:5, v/v) both containing 5 mM NH_4_HCO_2_ and 0.1% (v/v) formic acid. Separation was performed at 50°C with a flow rate of 0.3 ml/min using the following gradient: 0-15 min – 25 to 86 % B (curve 5), 15-21 min – 86 to 100 % B (curve 5), 21-34.5 min – 100 % B isocratic, 34.5-34.6 min – 100 to 25 % B (curve 5), followed by 8 min re-equilibration at 25 % B.

##### Mass Spectrometry

Reversed phase liquid chromatography was coupled on-line to a Q Exactive Plus Hybrid Quadrupole Orbitrap mass spectrometer (Thermo Fisher Scientific, Bremen, Germany) equipped with a HESI probe. Mass spectra were acquired in positive and negative modes with the following ESI parameters: sheath gas – 40L/min, auxiliary gas – 10L/min, sweep gas – 1L/min, spray voltage – 3.5kV (positive ion mode); −2.5kV (negative ion mode), capillary temperature – 250°C, S-lens RF level – 35 and aux gas heater temperature – 370°C. Data acquisition for lipid identification was performed in quality control samples by acquiring data in data dependent acquisition mode (DDA). DDA parameters featured a survey scan resolution of 140,000 (at *m/z* 200), AGC target 1e6 Maximum injection time 100 ms in a scan range of *m/z* 240-1200. Data dependent MS/MS scans were acquired with a resolution of 17,500, AGC target 1e5, Maximum injection time 60 ms, loop count 15, isolation window 1.2 *m/z* and stepped normalized collision energies of 10, 20 and 30 %. A data dependent MS/MS scan was triggered when an AGC target of 2e2 was reached followed by a Dynamic Exclusion for 10 s. All isotopes and charge states > 1 were excluded from fragmentation. All data was acquired in profile mode. For deep lipidome profiling, iterative exclusion was performed using the IE omics R package ^41^. This package generates a list for already fragmented precursors from a prior DDA run that can be excluded from subsequent DDA runs ensuring a higher number of unique MS/MS spectra for deep lipidome profiling. After the initial DDA analysis of a quality control sample, another quality control sample was measured but excluding all previously fragmentated precursor ions. Parameters for generating exclusion lists from previous runs were – RT window = 0.3; noiseCount = 15; MZWindow = 0.02 and MaxRT = 36 min. This workflow was repeated one more time to get a total of three consecutive DDA analyses of a quality control sample in positive ionization mode. Data for lipid quantification was acquired in *Full MS* mode with following parameters – scan resolution of 140,000 (at *m/z* 200), AGC target 1e6 Maximum injection time 100 ms in a scan range of *m/z* 240-1200.

##### Lipid Identification and quantification

Lipostar (version 1.0.6, Molecular Discovery, Hertfordshire, UK) equipped with *in house generated* structure database featuring fatty acids with no information on double bond regio- or stereoisomerism covering glycerolipid, glycerophospholipid, sphingolipid and sterol ester lipid classes. The raw files were imported directly with a *Sample MS Signal Filter Signal Threshold* = 1000 for MS and a *Sample MS/MS Signal Filter Signal Threshold* = 10. Automatic peak picking was performed with an *m/z tolerance* = 5 ppm,*chromatography filtering threshold* = 0.97,*MS filtering threshold* = 0.97, *Signal filtering threshold* = 0. Peaks smoothing was performed using the *Savitzky-Golay* smoothing algorithm with a *window size* = 3, *degree* = 2 and *multi-pass iterations* = 3. Isotopes were clustered using a *m/z tolerance* = 5 ppm, *RT tolerance =*0.25 min, *abundance Dev* = 40%, *max charge* = 1. Peak alignment between samples using an *m/z tolerance* = 5 ppm and an *RT tolerance* = 0.25 min. A gap filler with an *RT tolerance* = 0.05 min and a *signal filtering threshold* = 0 with an anti Spike filter was applied. For lipid identification, a “MS/MS only” filter was applied to keep only features with MS/MS spectra for identification. Triacylgylcerols, diacylglycerols and sterol esters were identified as [M+NH4]+ adducts. Phosphatidylcholines were identified as [M+HCOO] ^-^adducts. All other phospholipids were identified as [M-H]- adducts. Following parameters were used for lipid identification: 5ppm precursor ion mass tolerance and 20 ppm product ion mass tolerance. Automatic approval was performed to keep structures with quality of 3-4 stars. Identifications were refined using manual curation and Kendrick mass defect analysis and lipids that were not following these retention time rules were excluded as false positives. Quantification by integration of the extracted ion chromatograms of single lipid adducts was manually curated and adjusted. Identified lipids were normalized to peak areas of added internal standards to decrease analytical variation. For data representation, data was log10 transformed and autoscaled using metaboanalyst.ca^42^.

#### 5. Measurement of cardiolipin

##### Sample preparation

1 million cells/plate for 24 hr treatment or 2 million cells/plate for 2 hr treatment of selenite/vehicle cells were plated in 100 pi cell plates. After one day, the media was replaced with or without selenite. At 2 hr or 24 hr point incubation, media was removed, and the cells were harvested by scraping method within D-PBS (Thermo Fisher) as described in “sample preparation for lipidomics analysis”. The cell pellet was reconstituted with 400 µl D-PBS and snap frozen and sent out for the cardiolipin measurement.

##### Lipid extraction for mass spectrometry lipidomics

Mass spectrometry-based lipid analysis was carried out by Lipotype GmbH (Dresden, Germany) as described ^43^. Lipids were extracted utilizing a two-step chloroform/methanol procedure ^44^. Samples were spiked with a mixture internal lipid standards. After extraction, the organic phase was transferred to an infusion plate, then dried in a speed vacuum concentrator. The 1st step dry extract was re-suspended in 7.5 mM ammonium acetate in chloroform/methanol/propanol (1:2:4, V:V:V) and the 2nd step dry extract in 33% ethanol solution of methylamine in chloroform/methanol (0.003:5:1; V:V:V). All liquid handling steps were carried out using Hamilton Robotics STARlet robotic platform, with the Anti Droplet Control feature for organic solvents pipetting.

##### MS data acquisition

Samples were analyzed by direct infusion on a QExactive mass spectrometer (Thermo Scientific) that is equipped with a TriVersa NanoMate ion source (Advion Biosciences). Samples were analyzed in both positive and negative ion modes, with a resolution of Rm/z=200=280000 for MS and Rm/z=200=17500 for MSMS experiments, in a single acquisition. MSMS was triggered by an inclusion list that encompasses corresponding MS mass ranges scanned in 1 Da increments ^45^. Both MS and MSMS data were combined to monitor cardiolipin as deprotonated anions.

##### Data analysis and post-processing

Data were analyzed with an in-house developed lipid identification software based on LipidXplorer ^46, 47^. Data post-processing and normalization were carried out using an in-house developed data management system. Only lipid identifications with a signal-to-noise ratio greater than 5, and a signal intensity 5-fold higher than in corresponding blank samples were considered for further data analysis.

### Ubiquinone(CoQ) and Ubiquinol(CoQH_2_) analysis with UV spectrometry

Ubiquinone and ubiquinol have their unique pattern of the spectrum (Sup Fig 6c), so it is possible to verify ubiquinone to ubiquinol reduction by using UV spectrometry. 250 µl of 200 µM ubiquinone and ubiquinol solutions dissolved in EtOH were placed in the two wells each (facing each other) which is located in the center of a transparent 96 well plate. Then 100 µl of selenide gas-producing solution (described in the “Production of selenide gas” section) was placed in the center of a 96 well plate, and the plate cover was put on the plate bottom to let the ubiquinone/nol solution be exposed to selenide gas. After 5 min, the gas exposed ubiquinone and ubiquinol solutions in each well were pooled into each tube for the gas exposed ubiquinone and ubiquinol solution and centrifuged at 13,000rpm for 10 min to remove the red elemental selenium as elemental selenium interferes with the spectrum from ubiquinone/nol due to its own spectrum. The red precipitate is visible in the e-tube after centrifugation. The supernatant was added to the cuvette to measure absorbance values from 200nm to 600nm of gas exposed ubiquinone and ubiquinol solutions (Fig 3f,3g) with Nano Drop 1C (Thermofisher Scientific).

## Acknowledgments

We thank Arthur Mercurio, Cole Haynes, Nandhitha Uma Naresh, Sookyung Kim, and Olga Ponomarova for advice, assistance, and feedback. This work was supported by the Suh Kyungbae Foundation (SUHF) Young Investigator Award and a grant from the National Institutes of Health (R01CA269711) to D.K. N.L. and S.J.P. are recipients of Mogam Science Scholarship. J.B.S is funded by the Worcester Foundation Grant. This research was also supported by a grant from the National Institutes of Health (R01GM112948 to J.A.O.). J.A.O. is a Chan Zuckerberg Biohub investigator. P.L.G. is supported by fellowships from the Searle Scholars Program, the Rita Allen Foundation, and by grant DP2 OD027719­01 from the National Institutes of Health.

## Author contributions

N.L. and D.K. conceived the project; N.L. and D.K. designed the research. N.L. and S.J.P. performed most of the experiments with assistance from M.L., T.Y., M.B.D., Y.S., P.G., and J.S., and data analyses assistance and advice from T.Y.K., D.I.K., Y.S., M.L., J.O., and J.S. Cell viability related experiments were carried out by N.L. and S.J.P. ROS and other biochemical measurement experiments were carried out by N.L. and S.J.P. In vitro reactivity experiments were performed by N.L.. Immunocytochemistry was performed by M.B.D. Lipid profiling experiment was performed by M.L., N.L., and J.O. Polar metabolite profiling was performed by J.S. and N.L. Ubiquinone and ubiquinol analysis was performed by J.S., T.Y., N.L., S.J.P.,Y.S. Datamining, pathway analyses, and interpretation of metabolomics data were carried by M.L., J.O., T.Y. J.S., N.L., S.J.P., T.Y.K., and D.I.K. N.L., S.J.P., and D.K. wrote the manuscript, with consultation from all other authors.

## Materials & Correspondence

Materials and all raw data that support the findings of this study are available from the corresponding author upon request.

## Statistics and reproducibility

Results of the viability assay (Fig 1d,1i,4i and Sup Fig 2d,2e,3k,7c-7h,9a), mRNA quantification (Sup Fig 1a,1b), measurement of polar and lipid metabolites (Sup Fig 4b,4d,4e, Sup Fig 5h,5i), quantification of high ROS population (Sup Fig 4f,9e), iron population (Sup Fig 4h), superoxide radicals (Sup Fig 4h,4i), mitochondria depolarization (Sup Fig 9g), increased mitochondria population (Sup Fig 8h), Bid expression and localization to mitochondria (Sup Fig 9i,9j), selenide production capacity (Fig 2c,2e,2h, Sup Fig 3c,3d) and high lipid peroxidation population (Fig 1c,1g,4d,4g,4h, Sup Fig 2a,2b,2c,7h,8c,8j,9c), ubiquinol/none quantifications (Fig 3b,3g,3i,3m,3n,3o, Sup Fig 6d), total selenium measurement (Fig 2h,2i,3e), and total thiol measurement (Fig 2d,2e, Sup Fig 3h), were analyzed using Student’s t-test. P-values of less than 0.05 were defined as statistically significant, and graphical data marked with one (*), two (**) or three (***) asterisks represent p-values of < 0.05, < 0.01, and < 0.001, respectively. All error bars are standard deviation (SD). Most experiments were repeated at least three times with the following exceptions: Metabolite screen of polar and lipid metabolites (Sup Fig 4b,4d,4e,5a-5i), selenium ICP-MS quantification (Fig 2h,2i), LC-MS quantification of ubiquinol/ubiquinone after selenite supplementation in mouse and bacteria (Fig 3n,3o), and mRNA quantification of selenoproteins (Sup Fig 1a,1b) were carried out once in 3∼5 replicate samples. MS quantification of selenide-exposed ubiquinol/none (Fig 3g, Sup Fig 6d), and LC-MS quantification of ubiquinol/none quantification after selenite treatment in CTRL and SQOR KO cells (Fig 3l) were repeated twice. Figure Illustrations: Fig 1a, 1j, 2a, 4j, Sup Fig 6g, 6h were all created using Biorender.com.

---

**Supplementary Figure 1.**
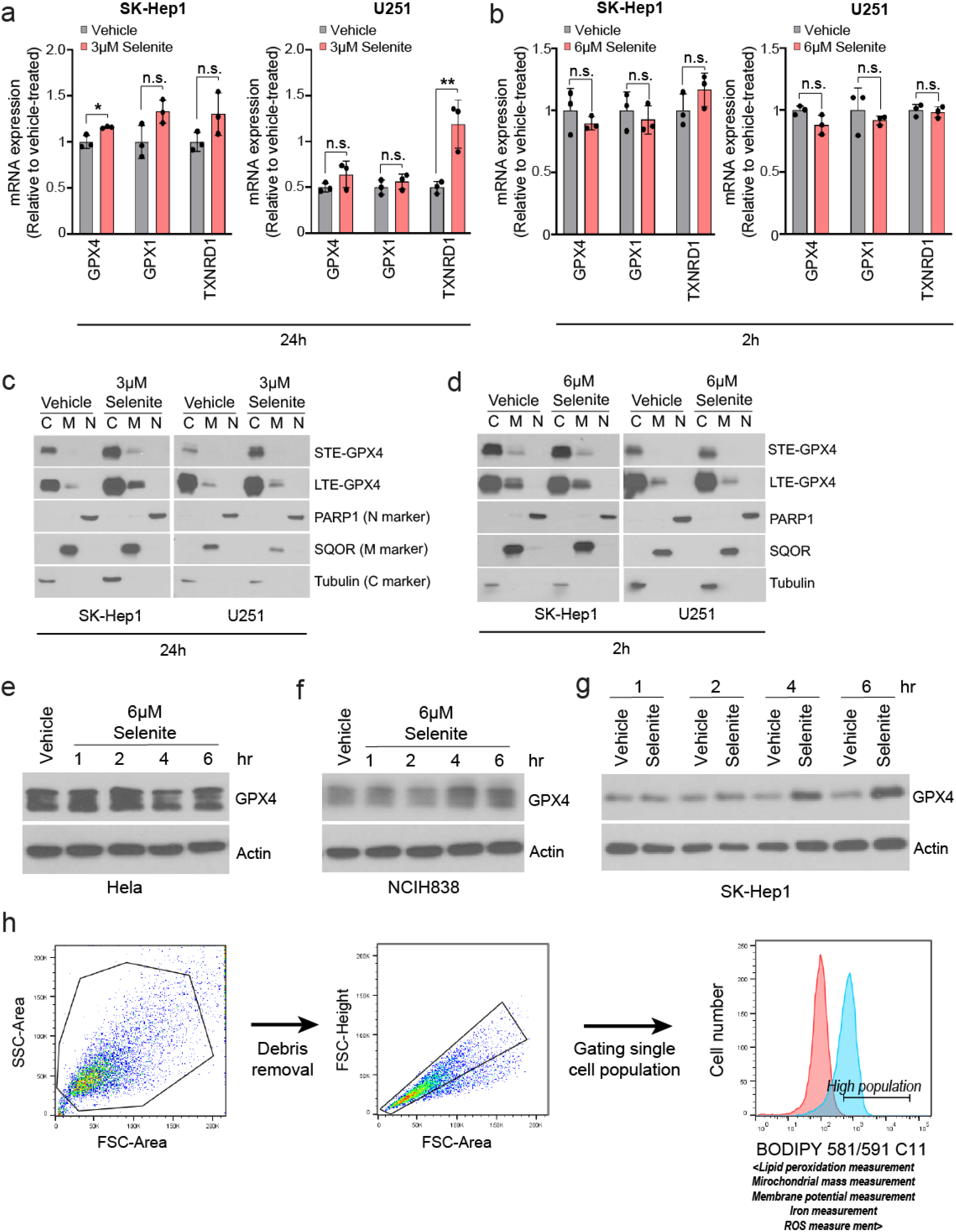
Selenium has an antiferroptotic effect that is independent of selenoprotein production, supplemental data 1. (a,b) Quantification of mRNA levels of selenoproteins in SK-Hep1 and U251 cells after selenite treatment for 24 hr (a) and 2 hr (b). (c,d) Immunoblots of GPX4 and organelle marker proteins in fractionated SK-Hep1 and U251 cells after vehicle or selenite treatment for 2 hr (d) and 24 hr (c). N, M, C, STE, and LTE represent nucleus, mitochondria, cytoplasm, short-term exposure, and long-term exposure, respectively. (e-g) Immunoblots of GPX4 in selenite treated Hela (e), NCIH838 (f), and SK-Hep1 (g) cells. (h) Representative flow cytometry data showing gating strategy for measurement of lipid peroxidation, iron, mitochondrial mass, and ROS. FSC-Area and SSC-Area gating strategy was used to eliminate cell debris. FSC-Height and FSC-Area subgating strategy was used to identify single cell population. The histogram of the single cell population was used for quantifying lipid peroxidation, iron, mitochondrial mass, or ROS. Data are mean ± S.D. from biological replicates (*n* = 3 for a,b) and were analyzed by two-tailed Student’s *t*-test. (a,b; * *P* < 0.05,\*\**P* < 0.01, *n.s.*, not significant).

**Supplementary Figure 2.**
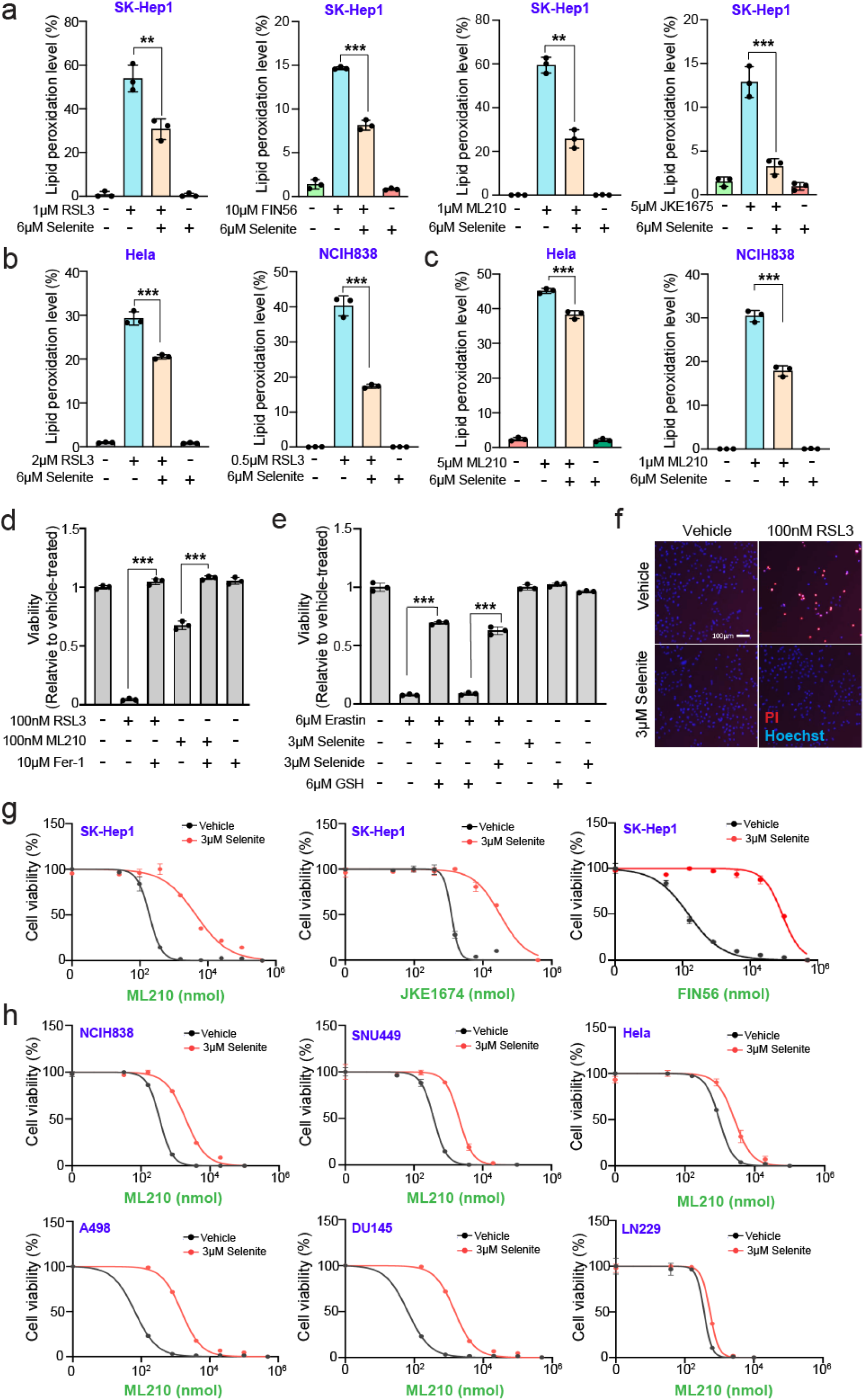
Selenium has an antiferroptotic effect that is independent of selenoprotein production, supplemental data 2. (a) Measurement of lipid peroxidation in SK-Hep1 cells treated with indicated doses of RSL3, FIN56, ML210, or JKE1675 with/without selenite treatment for indicated times. (b,c) Quantification of lipid peroxidation in RSL3 (b) or ML210 (c) treated Hela and NCIH838 cells with/without selenite for indicated times. (d) Viability of SK-Hep1 cells after treating with vehicle, RSL3, or ML210 with/without Ferrostatin for 24 hr. (e) Viability of SK-Hep1 cells after treating with vehicle or 6 µM erastin with/without 3 µM selenide or 3 µM selenite+6 µM L-GSH for 72 hr. (f) Fluorescence image of PI/Hoechst double stained SK-Hep1 cells treated with RSL3 and/or selenite for 24 h. Scale bar represents 100 µm. (g) ML210, JKE1675, and FIN56 dose response curve for vehicle or selenite treated SK-Hep1 cells. Cell viability was measured at 24 hr after treatment with the GPX4 inhibitors. (h) ML210 dose response curve for vehicle or selenite treated NCIH838, SNU449, Hela, A498, DU145, and LN229 cells. Data are mean ± S.D. from biological replicates (*n* = 3 for a-e,g,h) and were analyzed by two-tailed Student’s *t*-test. (a-e; \*\*\**P* < 0.001, \*\**P* < 0.01).

**Supplementary Figure 3.**
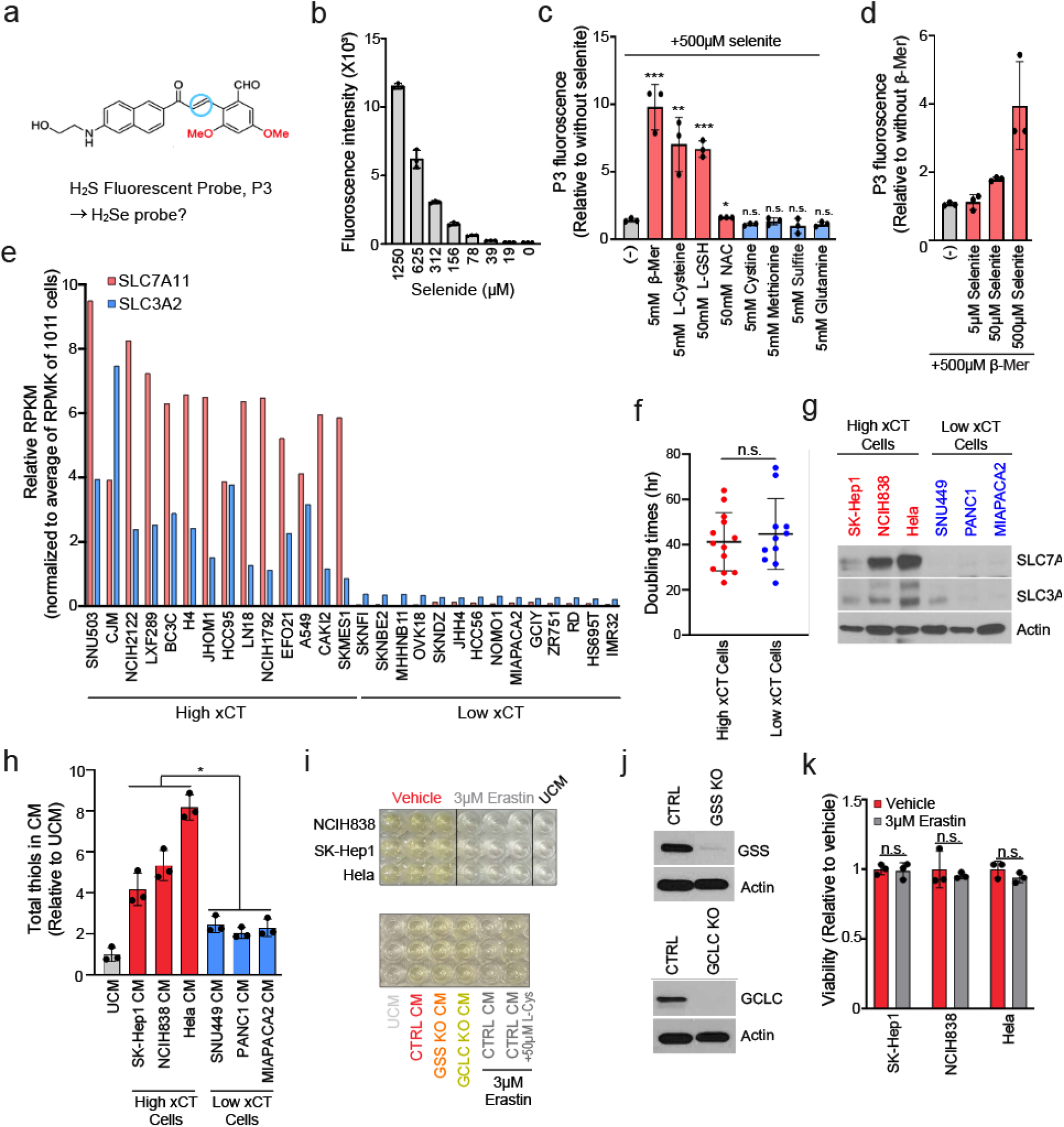
xCT mediated thiol formation induces selenite to selenide reduction. (a) Chemical structure of P3 probe originally developed for hydrogen sulfide detection^11^. (b) P3 probe-based selenide detection of different doses of selenide. Selenide results in increase of fluorescent intensity due to reaction with P3 probe. (c) Measurement of selenide levels using fluorescence based P3 probe after mixing with solutions of selenite and/or different metabolites as indicated. The selenite was added immediately after mixing P3 probe with thiol-containing metabolites (L-glutathione (L-GSH), L-Cysteine, β-mercaptoethanol (β-Mer), N-acetylcysteine), with negative controls of sulfur non thiol metabolite s (Cystine, Methionine, Sulfite), or non-sulfur, non-thiol metabolite Glutamate. Values are relative to fluorescent intensity of P3 probe mixed with each metabolite without selenite (=1.0). (d) Measuremen t of selenide immediately after mixing P3 probe with β-Mer and different doses of selenite. (e) RNA expressions from Cancer Cell Line Encyclopedia of the xCT subunits SLC7A11 and SLC3A2, used to designate xCT high and low cells. (f) Doubling time of high and low xCT cell lines represented in e. Doubling time information was collected from https://www.cellosaurus.org/. (g) Immunoblot of xCT subunits, SLC7A11 and SLC3A2 in three high xCT and low xCT cell lines. (h) Total thiol measurement of conditioned media from high xCT and low xCT cell lines conditioned for 24 hr. Each value is relative to that of the unconditioned medium (UCM), set to 1. (i) Wells containing Ellman’s solution for total thiol quantification in the conditioned media in Fig 2d,2e. (j) Immunoblot of GSS and GCLM from CTRL and GSS/GCLM KO SK-Hep1 cells. (k) Viability of high xCT cell lines treated with erastin for 24 hr. Values were normalized to that of vehicle treated cells (=1.0). Data are mean ± S.D. from biological replicates (*n* = 3 for c,d,h,k) and were analyzed by two-tailed Student’s *t*-test. (c,d,h,k; \**P* < 0.05,\*\**P* < 0.01,\*\*\**P* < 0.001, *n.s.*, not significant).

**Supplementary Figure 4.**
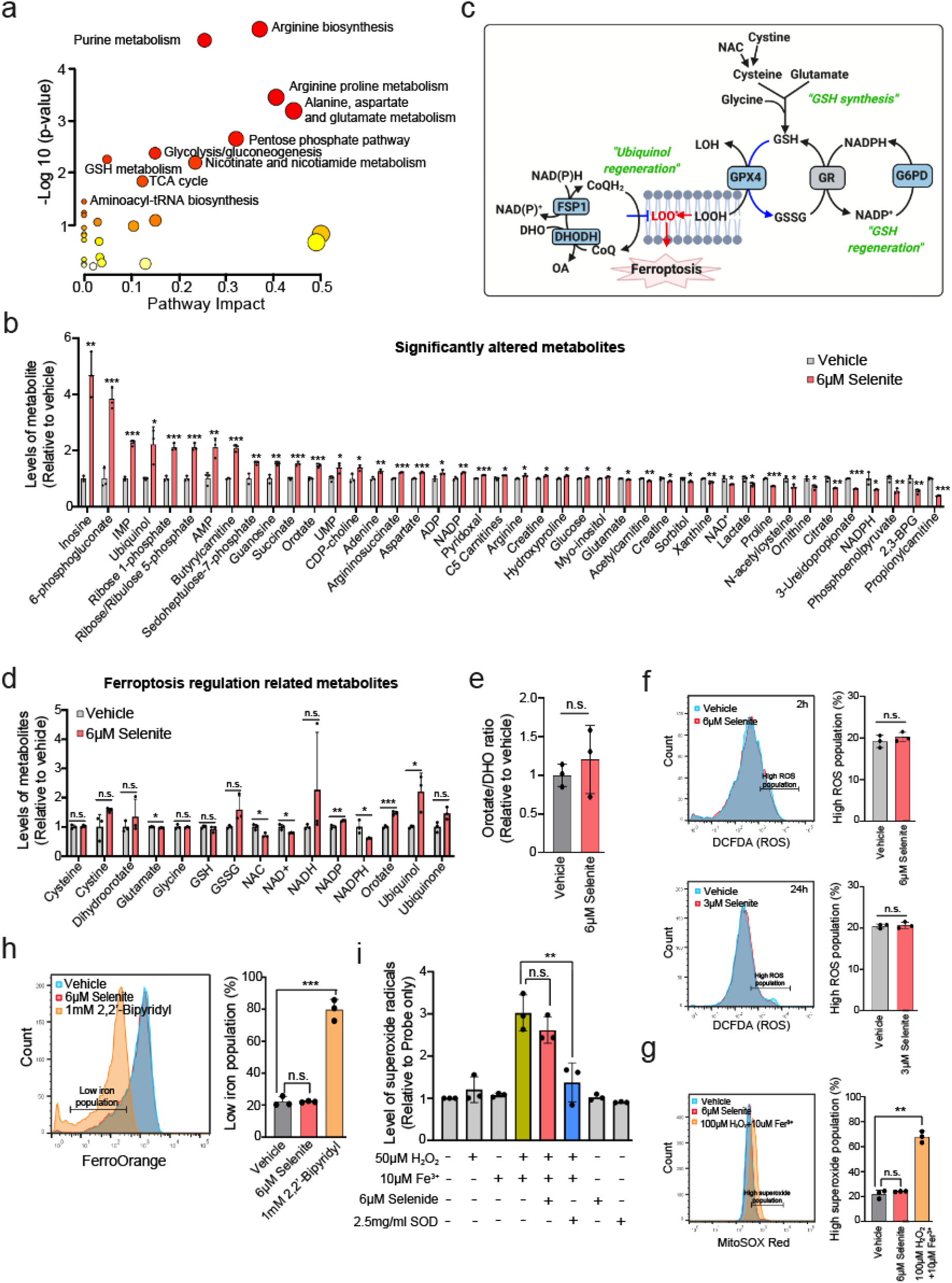
Effect of selenium on metabolite profile, ROS, superoxide, and iron levels. (a) Pathway analysis created with MetaboAnalyst 5.0 based on differential metabolites from vehicle and 6 µM selenite treated SK-Hep1 cells for 2 hr. A full list of metabolites is provided in Sup Table S2. (b) Relative levels metabolites that are significantly changed after 6 µM selenite treatment for 2 hr in SK-Hep1 cells. Values were normalized to that of vehicle treated cells (=1.0). (c) Schematic diagram of 15 key metabolites involved in the regulation of ferroptosis. (d) Levels of 15 key metabolites known to be involved in regulation of ferroptosis, measured in SK-Hep1 cells treated with vehicle or 6 µM Selenite for 2 hr. (e) The ratio of orotate and dihydroorotate (DHO) in SK-Hep1 cells treated with 6µM selenite for 2 hr, relative to vehicle treated cells (=1.0). (f) Quantification of intracellular reactive oxygen species (ROS) in SK-Hep1 cells after treatment of vehicle or 6 µM selenite for 2 hr or 3 µM selenite for 24 hr. Each bar graph (right) represents the high ROS population which is indicated as a bracketed bar in the histogram (left). (g) Measurement of mitochondrial superoxide radical in SK-Hep1 cells treated with vehicle, selenite, or combinations of H_2_O_2_ and Fe^3+^. (h) Measurement of intracellular iron in SK-Hep1 cells treated with vehicle, selenite, or iron chelator 2,2’-Bipyridyl used as a negative control. The bar graph next to the histogram indicates the percentage of the low iron population. (i) Superoxide levels combinations of H_2_O_2_ and Fe^3+^ which generates superoxide via Fenton reaction, demonstrating that selenide does not have radical trapping activity. Data are mean ± S.D. from biological replicates (*n* = 3 for a,b,d-i) and were analyzed by two-tailed Student’s *t*-test. (a,b,d-i; \**P* < 0.05,\*\**P* < 0.01,\*\*\**P* < 0.001, *n.s.*, not significant).

**Supplementary Figure 5.**
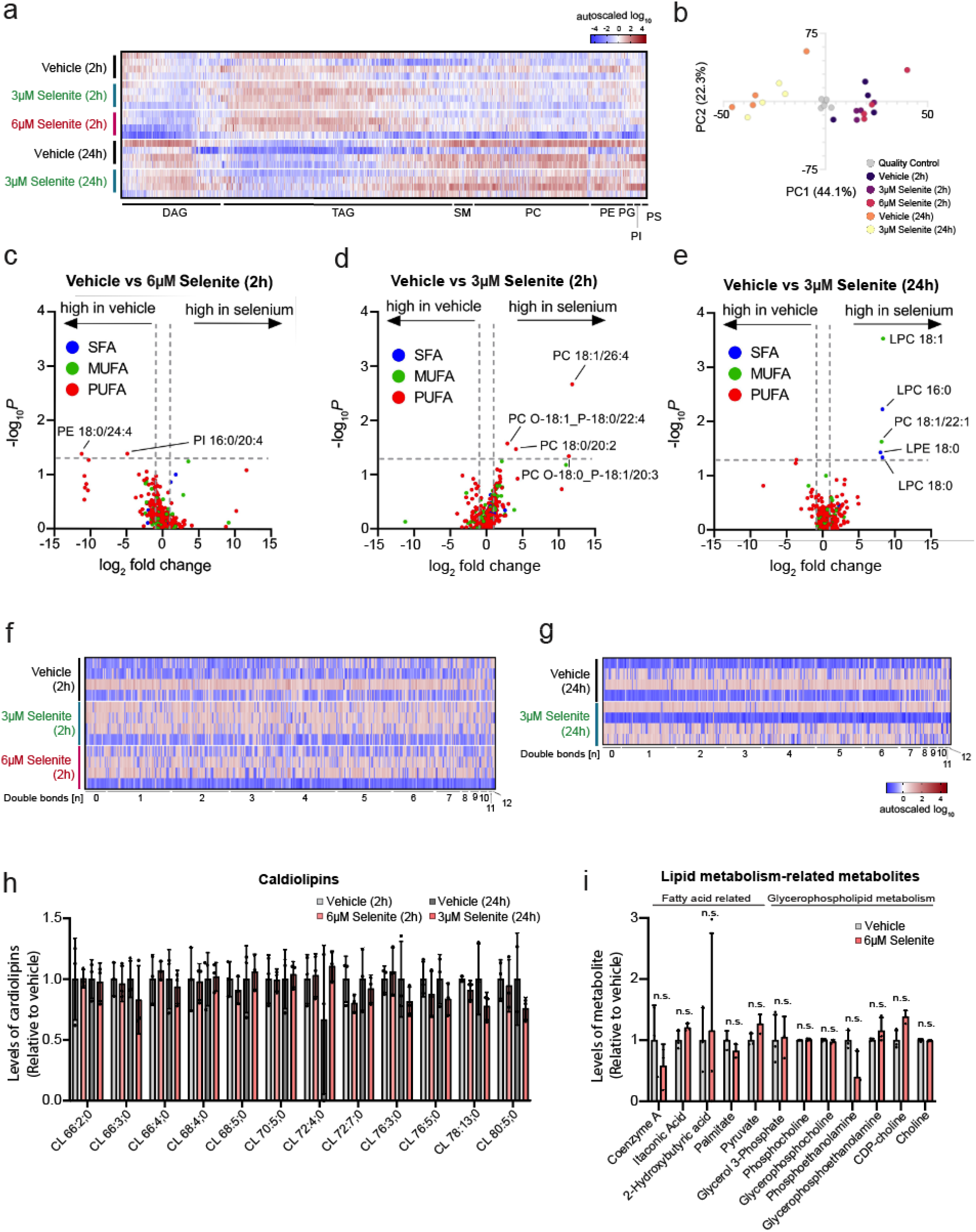
Effect of selenium on lipid profile. (a) Heatmap analysis of lipid profile from SK-Hep1 cells treated with selenite for 2 hr or 24 hr. The full lipid profiling data is provided in Sup Table S3. (b) Principal component analysis of lipid profile from SK-Hep1 cells treated with selenite for 2 hr o r 24 hr. (c-e) Scatter plot of lipid profiles of saturated fatty acids (SFA), monounsaturated fatty acids (MUFA), and polyunsaturated fatty acids (PUFA) from SK-Hep1 treated with 6 µM selenite for 2 hr (c), 3 µM selenite for 2 hr (d) and 3 µM selenite for 24 hr (e). (f,g) Heatmap analysis of lipid profile stratified by the number of double-bonds lipids from SK-Hep1 cells treated with selenite for (f) 2 hr and (h) 24 hr. h) Relative level of cardiolipin species in SK-Hep1 cells treated with 3 µM selenite or vehicle for 24 hr and 6 µM selenite or vehicle for 2 hr. Values were normalized to that of vehicle-treated cells (=1.0). (i) Relative levels of metabolites related to fatty acid and glycerophospholipid metabolism in SK-Hep1 cells after 6 µM selenite treatment for 2 hr. Values were normalized to that of vehicle-treated cells (=1.0). Data are mean ± S.D. from biological replicates (*n* = 3 for h,i, *n* = 4 for a-e) and were analyzed by two-tailed Student’s *t*-test. (h,i; *n.s*., not significant).

**Supplementary Figure 6.**
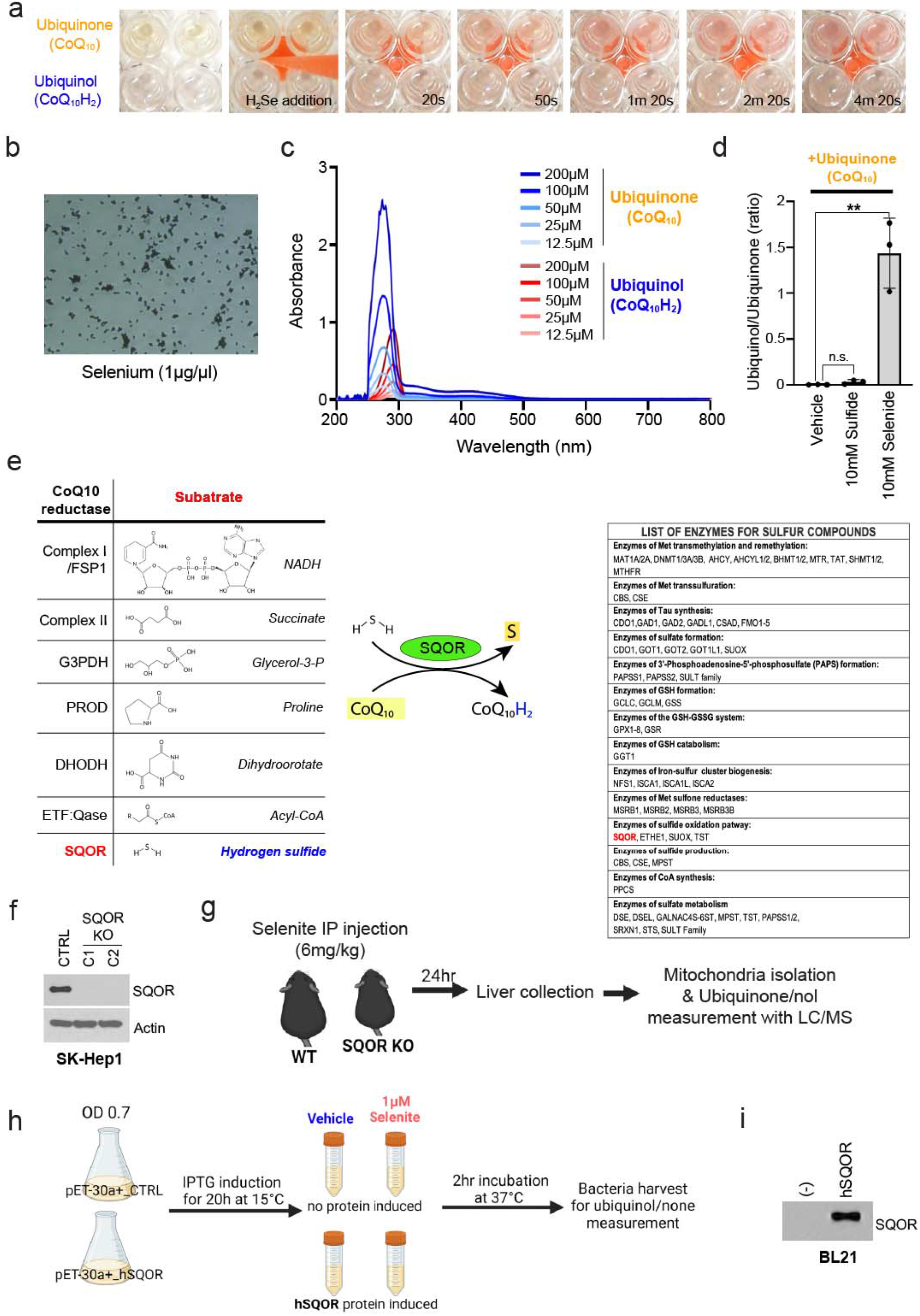
The selenium metabolite selenide reduces ubiquinone to ubiquinol via SQOR enzyme. (a) Time progression images of 96 well plate containing 200 µM ubiquinone/nol solutions before and after selenide gas exposure. Timepoints post gas exposure are indicated at the right bottom side of each image. (b) Brightfield images of solution containing 1 µg/µl selenium powder (Se). 100X magnification. (c) UV-vis spectrophotometer analysis of varying doses of ubiquinone and ubiquinol solutions. (d) Ubiquinol/none ratio in ubiquinone solution mixed with vehicle, 10 mM sulfide or 10 mM selenide solution. (e) Left, list of enzymes known to reduce ubiquinone to ubiquinol, and their known substrate. Middle, proposed chemical reaction of hydrogen sulfide by SQOR. Right, a list of enzymes that process sulfur-containing metabolites^48, 49^. (f) Immunoblot of SQOR in control and SQOR KO SK-Hep1 single clonal cells. (g) Schematic diagram of the experiment for testing the effect of selenite treatment on ubiquinol (CoQ_9_) production in the liver of wild type and SQOR KO mouse. (h) Schematic diagram of the experiment for testing the effect of selenite on ubiquinol (CoQ_8_) production in control and human SQOR-induced BL21 bacteria. (i) Immunoblot of SQOR protein in the hSQOR protein-induced bacteria. Data are mean ± S.D. from biological replicates (nV=V3) and were analyzed by two-tailed Student’s t-test. (d; \*\**P* < 0.01, *n.s.*, not significant).

**Supplementary Figure 7.**
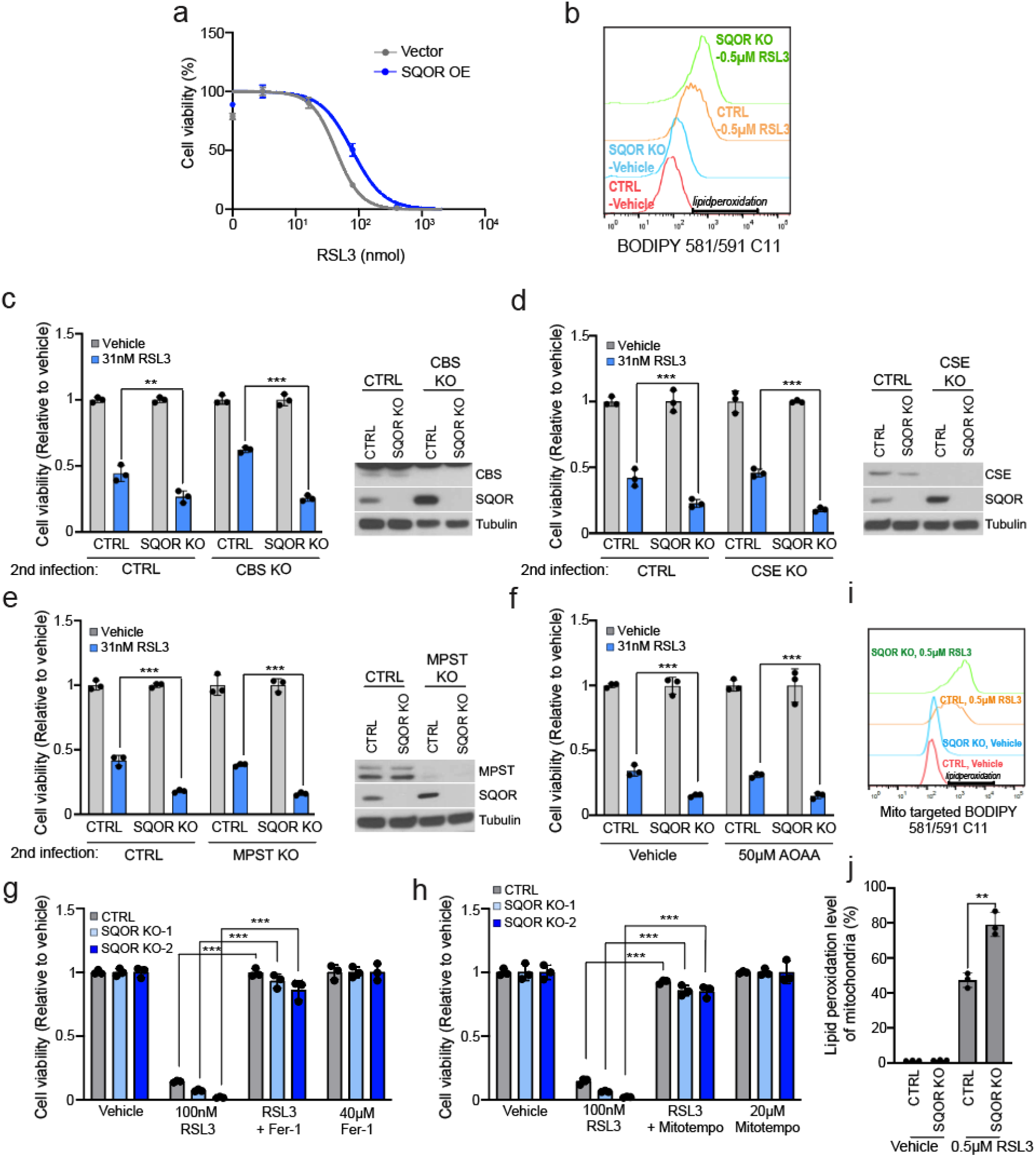
SQOR protects against ferroptosis. (a) Dose response curves for the GPX4 inhibitors RSL3 in LN229 glioma cells overexpressing blank vector or SQOR. Cell viability was measured at 24 hr after treatment with the GPX4 inhibitors. (b) Measurement of lipid peroxidation in CTRL and SQOR KO SK-Hep1 cells treated with vehicle or 0.5 µM RSL3 for 2 hr. Bracketed bar indicates the gating for peroxidized lipids. (c-e) Left, viability of hydrogen sulfide production impaired CTRL and SQOR KO SK-Hep1 cells after RSL3 treatment for 24 hr. Sulfide-producing enzyme genes, CBS (c), CSE (d), or MPST (e) were ablated by CRISPR-Cas9 in CTRL and SQOR KO SK-Hep1 cells. (f) AOAA, an inhibitor for sulfide - producing enzymes was treated in CTRL and SQOR KO SK-Hep1 cells with vehicle or RSL3 for 24 hr. (c-e) Right, immunoblot of (c) CBS, (d) CSE, and (e) MPST of CTRL and SQOR KO SK-Hep1 cells, which were subjected to additional KO with CTRL or CBS/CSE/MPST. (g) Viability of CTRL and SQOR KO SK-Hep1 cells treated with RSL3 and/or Ferrostatin-1 (Fer-1) for 24 hr. (h) Viability of CTRL and SQOR KO SK-Hep1 cells treated with RSL3 and/or Mitotempo for 24 hr. (i) Measurement of mitochondrial lipid peroxidation in CTRL and SQOR KO SK-Hep1 cells treated with vehicle or RSL3 for 2 hr. Bracketed bar indicates the gating for quantifying mitochondrial lipid peroxidation. (j) Quantification of peroxidized lipids from panel g. Data are mean ± S.D. from biological replicates (nV=V3) and were analysed by two-tailed Student’s *t*-test. (c-f,h-j; \*\**P* < 0.01,\*\*\**P* < 0.001).

**Supplementary Figure 8.**
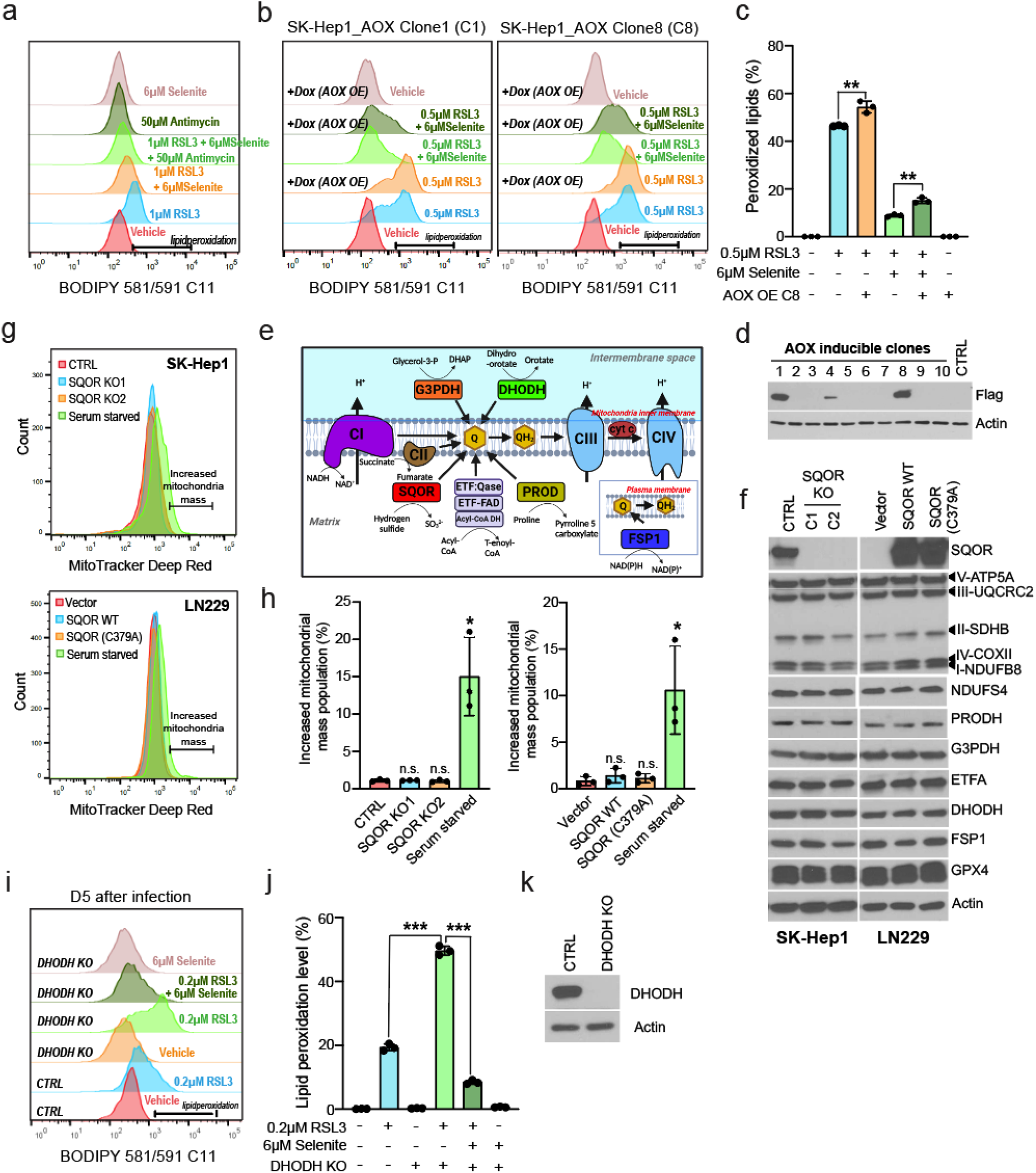
SQOR protects against ferroptosis, supplemental data. (a) BODIPY dye measurement of lipid peroxidation in RSL3 or/and selenite-treated SK-Hep1 cells with/without Antimycin, concomitantly for 2 hr. Bracketed bar indicates the gating for lipid peroxidation. (b) BODIPY dye measurement of lipid peroxidation levels of SK-Hep1 cells with/without induced AOX overexpression in two independent clonal cell lines, as indicated in panel d, treated with/without RSL3 for 2 hr. (c) Quantification of lipid peroxidation level from AOX clone 8 in panel b. (d) Immunoblot of flag-tagged AOX protein. 10 single clones obtained after viral infection containing AOX expression construct were treated with doxycycline for 48h and collected samples for confirming the expression of AOX protein. Clone 1 and 8 were used for panels b and c. (e) Schematic diagram of enzymes involved in ubiquinone reduction to ubiquinol. The substrate and product of the enzymes are depicted. (f) Immunoblots of components of the electron transport chain and CoQ10 oxidoreductases, which affect the levels of ubiquinol, and key ferroptosis regulators in SQOR KO SK-Hep1 cells and SQOR OE/SOQR mut OE LN229 cells. ATP5A, ATP synthase lipid-binding protein; UQCRC2, Ubiquinol-cytochrome c reductase Core Protein 2; SDHB, Succinate dehydrogenase complex iron sulfur subunit B; COXII, Cytochrome c oxidase subunit 2; NDUFB8, NADH:Ubiquinone oxidoreductase subunit B8; NDUFS4, NADH:Ubiquinone oxidoreductase subunit S4; PRODH, proline dehydrogenase; G3PDH, Glyceraldehyde-3-phosphate dehydrogenase; ETFA, Electron transfer flavoprotein subunit alpha; DHODH, Dihydroorotate dehydrogenase; FSP1, Ferroptosis Suppressor Protein 1; GPX4, Glutathione peroxidase 4. (g) Mitotracker dye quantification of mitochondria in CTRL or SQOR KO SK-Hep1 cells and Vector, SQOR OE, or SQOR mut OE LN229 cells. Bracketed bar indicates the gating for increased mitochondria population. Cells incubated with 1% serum for 36 hr were used as a control. (h) Quantification of high mitochondria population from in panel g. (i) BODIPY dye measurement of lipid peroxidation in CTRL and DHODH KO SK-Hep1 cells treated with RSL3 and/or selenite for 2 hr. SK-Hep1 cells at 5 days after infection of virus containing guide RNA targeting DHODH were used. Bracketed bar indicates the gating for lipid peroxidation. (j) Quantification of lipid peroxidation from panel i. (k) Immunoblot of DHODH in CTRL and DHODH KO SK-Hep1 cells. Data are mean ± S.D. from biological replicates (*n* = 3 for c,h,j) and were analyzed by two-tailed Student’s t-test. (c,h,j; * *P* < 0.05,\*\**P*<0.01,\*\*\**P* < 0.001, *n.s.*, not significant).

**Supplementary Figure 9.**
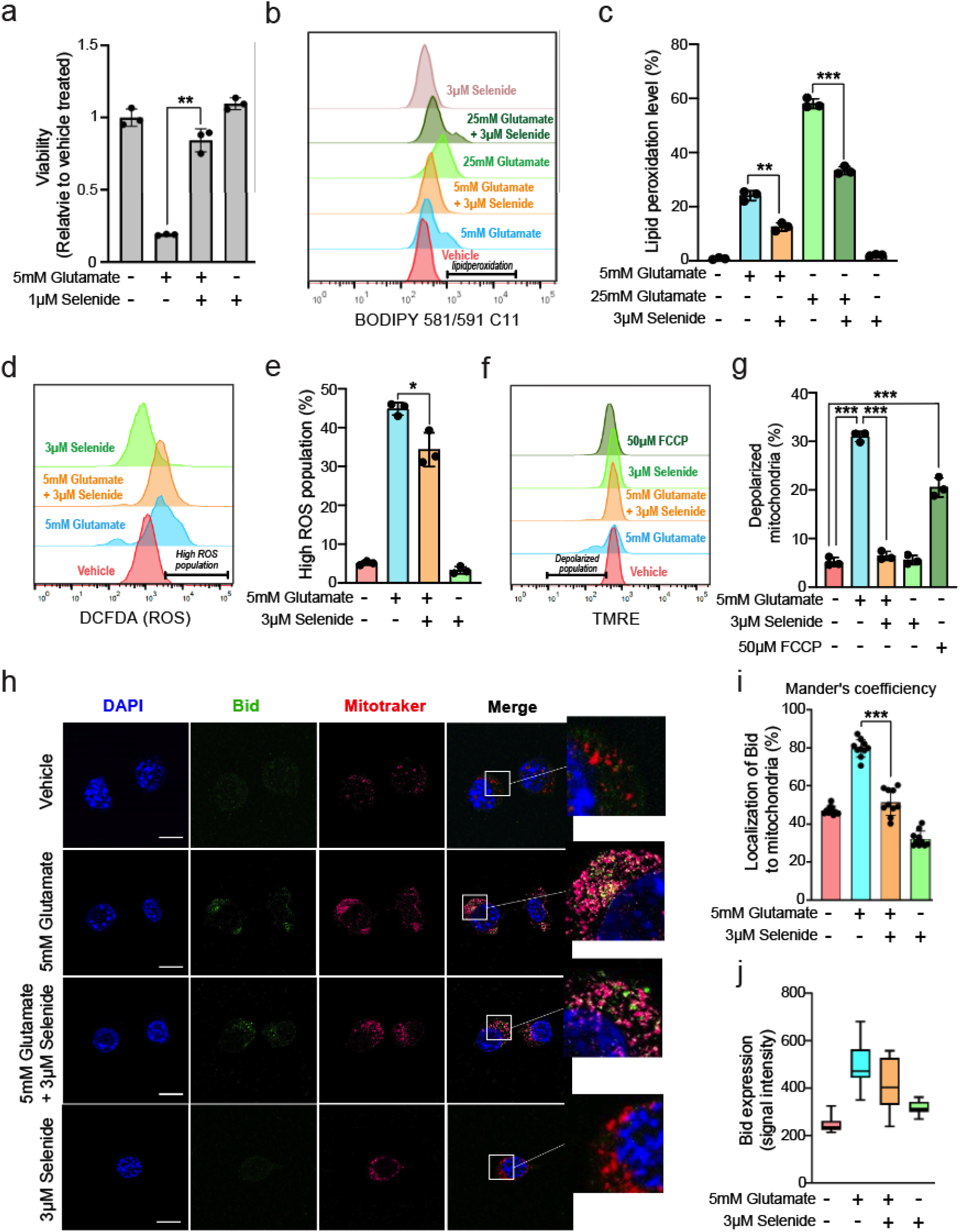
Selenide protects hippocampal neurons from glutamate induced ROS, lipid peroxidation and membrane depolarization. (a) Viability of mouse hippocampal neuronal HT29 cell s after 5 mM glutamate treatment for 24 hr with/without selenide, relative to vehicle treated cells (=1.0). (b) BODIPY dye measurement of lipid peroxidation in HT29 cells treated with 5 or 25 mM glutamate and/or 3 µM selenide for 12 hr. Bracketed bar indicates the gating for peroxidized lipids. (c) Quantification of lipid peroxidation from panel b. (d) DCFDA dye measurement of ROS in HT29 cells treated with 5 mM glutamate and/or 3 µM selenide for 12 hr. Bracketed bar indicates the gating for high ROS population. (e) Quantification of high ROS population from panel d. (f) TMRE dye measurement of mitochondrial membrane potential in HT29 cells treated with 50 µM FCCP or 5 mM glutamate and/or 3 µM selenide for 12 hr. The uncoupler FCCP was used as a control. Bracketed bar indicates the gating for the depolarized population. (g) Quantification of the depolarized population from panel f. (h) Immunocytochemistry of Bid protein, mitochondria, and nucleus in HT29 cells treated with 5 mM glutamate and/or 3 µM selenide for 12 hr. Scale bar indicates 20 µm. (i) Quantification of Bid protein localized to mitochondria from panel h. (j) Quantification of Bid protein expression from panel h. Data are mean ± S.D. from biological replicates (*n* = 3 for a,c,e,g; n = 10 for i,j) and were analyzed by two-tailed Student’s t-test. (a,c,e,g,i; \**P* < 0.05,\*\**P* < 0.01,\*\*\**P* < 0.001, *n.s.*, not significant).

